# A cross-frequency architecture links biological and behavioral periodicities

**DOI:** 10.64898/2026.07.03.736377

**Authors:** Antonio Criscuolo, Tongxin Liu, Michael Schwartze, Sonja A. Kotz

## Abstract

Spontaneous behaviors, e.g., walking and speaking, are thought to rely on an internal *sense* of time that provides a scaffold for precise temporal coordination. Yet endogenous rates of many behavioral processes often diverge from arbitrarily defined unit of *objective* time, raising a fundamental question: what temporal reference frame coordinates behavior?

Using high temporal resolution multimodal recordings, we show that endogenous body-brain periodicities – including pupil fluctuations, saccadic eye movements, respiration, cardiac activity and neural oscillations - as well as spontaneous behaviors such as tapping, walking, and speaking, form a cross-frequency architecture organized as an arithmetic progression in a natural logarithmic frequency space. Successive rates are spaced by about φ (golden ratio ≈ 1.618 ^1^), while the ratio between rhythm classes are captured by *e* (*e* ≈ 2.72 ^2^). The absence of power law scaling and harmonic anchoring to individual heart rates excluded the possibility that the frequency architecture scales to, and depends uniquely on, a single pacemaker. Rather, the irrational scaling suggests each rate to be principally independent and maximally de-coupled from others, yet capable to dynamically engage in cross-frequency interactions. We propose that the body-brain frequency architecture may provide an emergent temporal reference frame to scaffold perception and action coordination.

## Introduction

“*The second is the duration of 9192631770 periods of the radiation corresponding to the transition between the two hyperfine levels of the ground state of the caesium 133 atom.*”

- 13th General Conference on Weights and Measures in 1967.

Inspired by James Clerk Maxwell, the International Atomic Time (IAT) connects the world into a unique *sense* of time, which discretizes the continuous flow of time utilizing the frequency of electromagnetic radiation of the caesium 133 atom. This socially *agreed-upon, arbitrarily defined, ‘physical’* measure of time ^2^ anchors daily life routines across the globe, coordinates spatial navigation and transportation systems, structures work shifts, and countless other activities. As humans grow up and become accustomed to utilizing a common reference frame (IAT) for estimating the flow of time, one may expect *physical time* to be leveraged to coordinate perceptual processing and movement control. In this view, if everyone *ticks* according to a common *clock* in the scale of seconds, there should be minimal inter-individual variation in time estimation performance (i.e., estimating temporal intervals between sensory events (i.e., Δ*_t_* = A*_t1_*-B*_t2_*), preferred perceptual tempo ^3–5^, and in the spontaneous pacing of movements (e.g., spontaneous tapping rhythm). In fact, absolute *physical* time (*Chronos*) should leave no space to *subjective* time (*Kairos*). However, when asked to move at a spontaneous pace, people rarely tap or walk at a second (1Hz) rate: spontaneous motor tempo (SMT) diverges from the second, as well as from its (sub)harmonics ^5^, ultimately ranging between ∼1.3 – 1.8Hz ^4,6–9^. This observation is not trivial: the act of spontaneously pacing behavior (e.g., tapping) may represent individual strategies to *discretize* the continuous flow of time into consecutive temporal intervals. In this view, the internal sense of time is embodied, and emerges via the *spatialization* of the arrow of time ^10,11^. That is why SMT has been traditionally regarded as an opportunity to investigate subjective representation of time, and more generally endogenous time processing ^6^.

If SMT diverges from the arbitrary units defined by *artificial* clocks, what is the internal reference frame utilized to coordinate behavior in time? Secondly, intra- and inter-individual variations in SMT ^6,12^ make us wonder: if the *sense of time* is not fixed, what determines its *fluctuations* within and across individuals? These questions are core to a fundamental understanding of general cognition in living species: as in natural sciences time is used to characterize motion, dynamics and to derive predictions about future events, our brain utilizes time not only for coordinating action cycles, but more generally to optimize information processing, perception and adaptive behaviors ^13–16^.

In a recent opinion piece, Buzsáki ^2^ proposed the internal representation of time to be linked to a hierarchy of body-brain rhythms. The core tenets of the proposed model are the following: (H1) the hierarchy of body-brain rhythms forms a linear progression on a natural logarithmic scale; (H2) the variability within each class grows with its mean frequency, in line with the Weber–Fechner principle that governs many biological and perceptual magnitudes (e.g., space, time and memory processing) ^16–18;^ (H3) changes in body-brain states induce fluctuations in the internal representation of time.

### Natural logarithmic organization

In the brain, characteristic frequency bands (from slow oscillations through delta, theta, gamma bands and fast ripples) are approximately equidistant on a natural-logarithmic axis, so that neighboring classes differ by a roughly constant ratio, *e*: 3.14 ^19,20^. The log-spacing of frequencies allows for high efficiency and robustness ^19^, ensuring that harmonic interference is minimized – i.e., if frequencies were linearly spaced, the harmonics of lower frequencies would constantly disrupt higher-frequency bands. The natural logarithmic distribution emerges from the temporal nesting of multiplexed processes (e.g., expressed in the form of cross-frequency phase–amplitude coupling) in neural oscillatory activity ^19,21,22^, whereby the phase of a slower rhythm modulates the amplitude of faster ones, allowing short events to be grouped into longer units. By extending these principles from the brain to the body, autonomic, respiratory and cardiac rhythms are proposed to expand the same logarithmic series towards slower timescales, and to be coupled to brain oscillations through ascending interoceptive pathways ^23–25^.

Pairing recurring bodily events with neural oscillations would provide a continuous calibration mechanism to ground the subjective representation of time. This framework expands psychological ^13,17,26–28^ and neural ^16,29–32^ models of temporal cognition, mapping the wide span of experienced time onto a *dynamic* cross-scale frequency architecture offered by biological periodicities in the body and the brain ^33,34^. This view embraces earlier conceptualizations on the influence of mind-body states on the perception and estimation of time put forward by Fraisse ^15^ and Church ^28^. Unlike *Chronos, Kairos* allows the internal sense of time to speed up or slow down, stretching and contracting experienced durations in function of changes in bodily states ^35^.

### The alternative: golden-mean spacing for body-brain coupling

While agreeing on the existence of a body-brain frequency architecture, Klimesch ^1,36^ proposed a complementary, more fine-grained architecture. Its central claim concerns how neighboring frequencies should be spaced to balance two competing needs: rhythms must be able to couple transiently when information has to be exchanged; yet, they must not lock to each other permanently, which would abolish their independence. The ratio that best satisfies both constraints is identified in the ‘golden ratio’ (φ ≈ 1.618). Because φ is the irrational number least well approximated by ratios of integers, rhythms separated by φ never settle into a stable phase relationship (minimal spurious synchronization), whereas small frequency shifts can bring them transiently into register ^37^ . So far, this framework was confirmed in resting EEG frequencies, where center frequencies were found to be organized as a binary (doubling) hierarchy. Klimesch proposed to extend this organizational principle to include bodily rhythms in the architecture. This framework generates two further hypotheses: (H4) successive body-brain-behavioral (BBB) rhythms are separated by a ratio close to φ; and (H5) individual rhythms are anchored to a bodily reference rhythm, the heart rate (HR), at harmonic (octave) positions. In turn, individuals with a faster HR would also show proportionally faster rhythms elsewhere.

### The current study

The two frameworks are not mutually exclusive: they both reinforce the notion that the body and the brain are complex and highly interconnected dynamic systems ^34,38–41^, but propose different scaling laws to coordinate rhythms across modalities. Importantly, while Buzsáki’s hierarchy describes how broad classes of rhythms partition the frequency axis, Klimesch’s golden mean describes how neighboring rhythms are arranged, and whether this arrangement is anchored in individual physiology. Testing them jointly therefore requires analyses at three levels: rhythm classes, individual modalities, and individual participants. Next to these core analyses, we assessed the relationship between bodily and spontaneous behavioral rhythms. We reasoned that two scenarios are possible (H6): either behavioral rates are random and not associated with bodily rhythms; or they would fall within the rhythm classes predicted by the body-brain hierarchy progression on the frequency axis. In turn, the body-behavior relationship could take two forms with different implications: under rigid anchoring, a person’s behavioral rates would scale with their own bodily rhythms, as assumed by single-pacemaker models of timing ^17,28^. Alternatively, under structural anchoring, behavior would occupy the same frequency classes as body–brain rhythms at the population level, and would be able to vary across individuals within each defined class.

In this study, we addressed the six hypotheses listed above by recording multimodal body-brain-behavioral data at rest, in perception and action. Combined evidence from pupillometry, saccadic eye movement, respiratory, cardiac, neural activity, and spontaneous behavior confirms the existence of a cross-scale frequency architecture linking biological and behavioral periodicities on a natural logarithmic scale.

## Results

By combining high-temporal resolution multimodal data (Fig. 1), in this work we aimed at (i) characterizing individual endogenous rates in the body, brain and behavior (BBB) and (ii) assessing the statistical relationship between rhythms across modalities. Below, we provide results for the two primary aims and then introduce additional exploratory analyses on the modulatory role of interoceptive abilities (‘Exploratory analyses – Part I’) and the link between individual heart rates and cardiac cycle with other body-brain-behavioral rates (‘Part II’).

**Figure 1.**
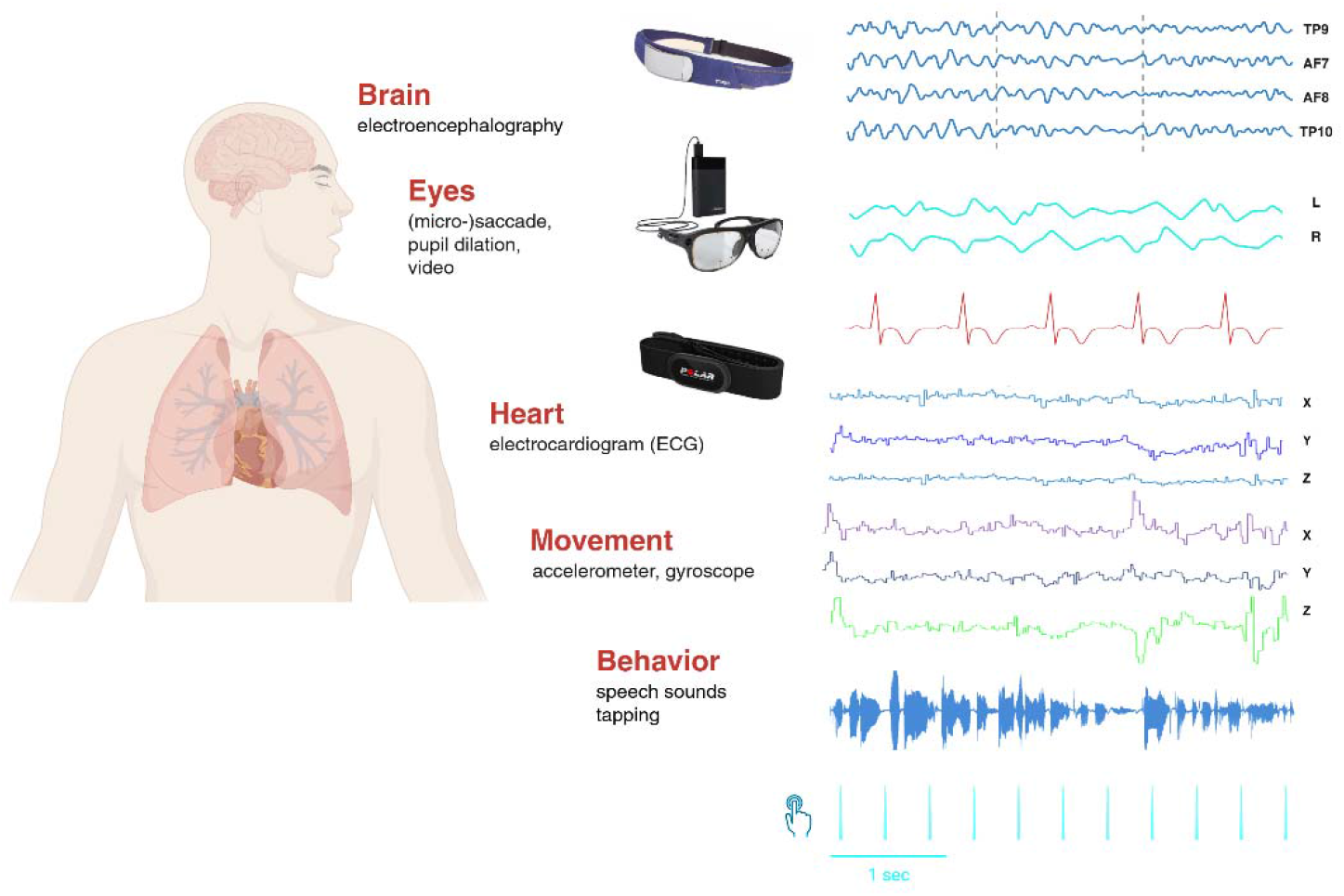
Experimental setup and multimodal body-brain-behavioral data acquisition. We recorded brain activity with a wearable EEG system (MUSE) with 4 channel locations: TP9 and TP10, AF7 and AF8. The headband further collected motion data via 3D accelerometer and gyroscope sensors. We also registered eye movements and pupil dilation via Tobii smart wearable glasses. A Polar H10 chest band was employed to record ECG activity as well as 3D accelerometer data. Finally, an Android app integrated signals from the MUSE and Polar bands, further recording audio signals and taps on the screen.

### BBB rhythms

We quantified individual endogenous rates in the body (e.g., see Suppl. Fig. 2A-B for ECG and respiration; 2D for pupil dilation) and the brain (see Suppl. Fig. 2C for IAF), as well as individual rates in spontaneous behavior (see Suppl. Fig. 1 for gait cycle; Suppl. Fig. 3 for speaking; Suppl. Fig. 4 for rhythmicity analyses on tapping and walking rates).

Discrete and continuous rate analyses (see details in Methods) demonstrated a predominant rhythm for all participants and in every modality, except for saccadic eye movements. Saccades during walking did not display periodic timing and are consequently treated as ‘characteristic’ median rate, but not as an *endogenous* rhythm.

Multimodal BBB rhythms are provided in Fig. 2. In order we display inter-individual rates (Fig. 2A and Fig. 2B) in pupil dilation (PDil = .2Hz), breathing (BR = .3Hz), tapping (TR = 1.4Hz), heart (HR = 1.4Hz)

**Figure 2.**
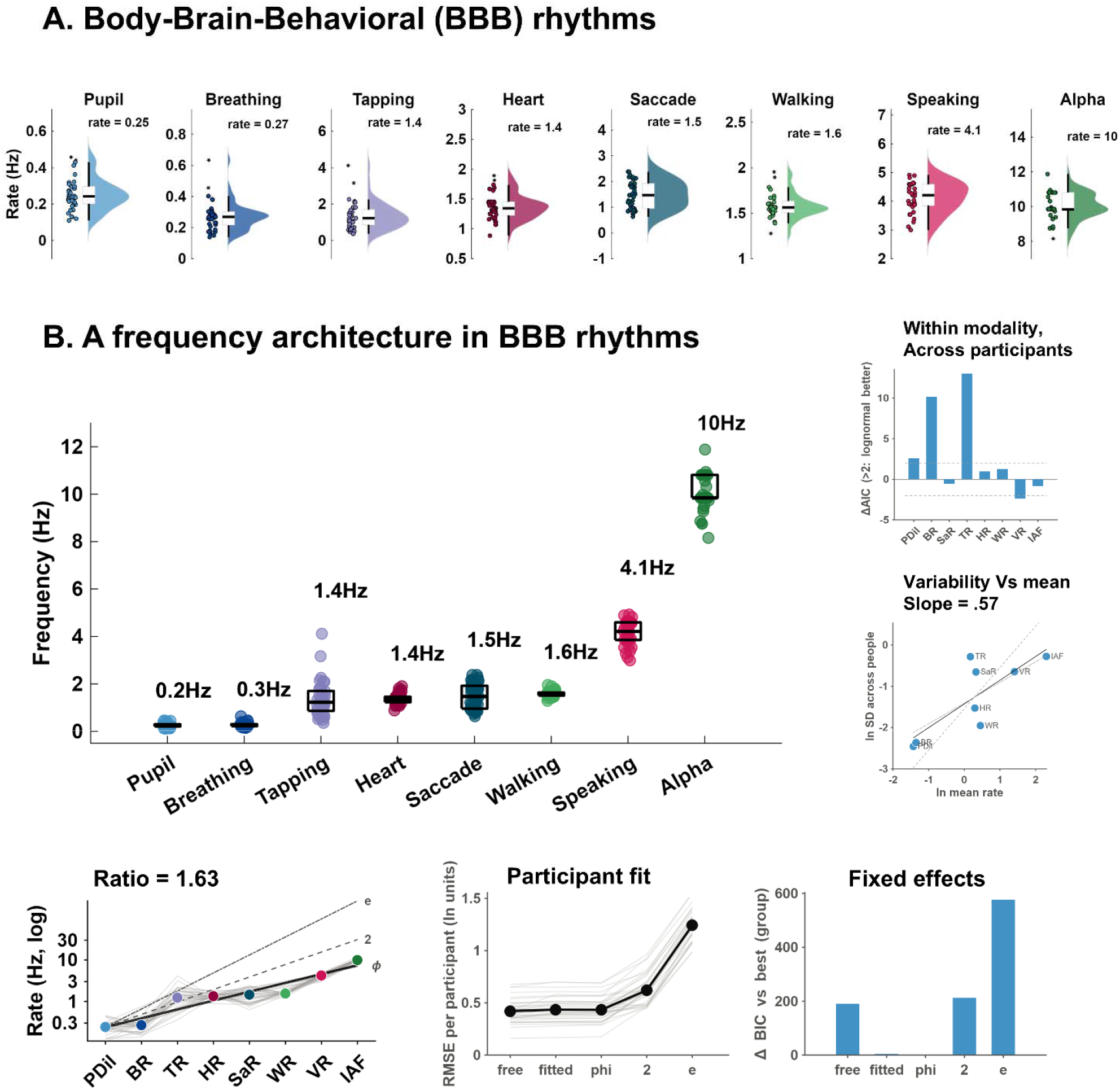
Body-Brain-Behavioral Rhythms. A: We characterized endogenous rates in, in order: pupil dilation (PDil), breathing (BR), tapping (TR), heart (HR), saccadic eye movement (Saccade), walking (WR), speaking (VR) rates as well as individual alpha frequency (IAF). Plots show the distribution of rates across 44 participants B: endogenous body-brain-behavioral rates are disposed on a frequency axis. On the right, top panel: Shape of inter-individual distributions within each modality. Bars show the difference in Akaike information criterion between normal and lognormal fits of individual rates (ΔAIC = AIC_normal − AIC_lognor al). Positive values favor a lognormal distribution (right-skewed, proportional individual differences, consistent with multiplicative generative processes); negative values favor a normal distribution (symmetric differences around a characteristic rate, consistent with additive perturbations of a set point). Dashed lines mark ΔAIC = ±2. One observation per participant per modality. Below, Scaling of inter-individual variability with frequency. Each dot represents one modality: log standard deviation of individual rates across participants against log geometric-mean rate. The solid line indicates the least-squares fit (slope = 0.57). The dashed line shows slope 1, i.e. constant coefficient of variation; the dotted line shows slope 0.5, i.e. variance proportional to the mean. Points above the fit vary more between individuals than expected from their frequency, points below vary less. Bottom row, on the left: Rates of eight rhythms on a logarithmic axis, ordered by group median. We calculated the log-ratio between adjacent rhythms and compared five descriptions of each participant’s rates across modalities: an individual model (‘free’); a common empirical model (leave-one-participant-out; ‘fitted’); three fixed ladders with ratios set to φ (phi: 1.618), 2 (harmonic) and ‘e’ (2.41). Each grey line represents one participant. Colored dots display the group mean per modality. black line: average constant-ratio fit (ratio per step 1.63). Bottom, middle plot: we computed the root-mean-square deviation of each participant’s log-rates from five ladders: individual ratio (free), common ratio estimated from the other participants (fitted), and fixed ratios φ, 2 and e. Grey: participants; black: medians. Bottom right: group evidence for each ladder: Bayesian information criterion (BIC) relative to the best model (lower is better; differences > 10 are very strong evidence).

saccadic (SaR = 1.5Hz), walking (WR = 1.6Hz), speaking (VR = 4.1Hz) rate and individual alpha frequency peak (IAF = 10Hz). After obtaining individual median rates, single-participant rates were sorted in ascending order in Fig. 2B.

### Distribution of individual rates within and across modalities

The shape of the inter-individual distribution differed between rhythms (Fig. 2B, bottom left). A lognormal model was clearly preferred for tapping rate (ΔAIC = [13.1]) and breathing rate ([10.2]), and modestly for pupil-oscillation rate ([2.6]). These rhythms thus showed right-skewed, proportional individual differences, consistent with multiplicative underlying processes. For heart rate ([1.0]), walking cadence ([1.3]) and individual alpha frequency ([−0.8]), the two models fitted equally well. Inter-individual variability was small for these rhythms (CV = [0.16], [0.09] and [0.08], respectively), a regime in which normal and lognormal distributions are practically indistinguishable. Their rates are therefore best described as narrowly distributed around a characteristic value. Syllabic rate was slightly better fitted by a normal distribution ([−2.4]; CV = [0.13]), indicating symmetric individual differences around a characteristic speech rate.

Across modalities, the inter-individual SD increased with mean rate with a slope of [0.57] (95% CI [.49 .67]; Fig. 2B, bottom middle panel). This is below the slope of 1 expected for a constant CV. Faster rhythms were thus relatively less variable between individuals than slower ones. The residuals around this relation suggested a grouping by the degree of constraint rather than by frequency. Self-paced tapping and the slow autonomic rhythms (breathing, pupil) varied most between individuals (CV ≈ [0.64], [0.35], [0.35]). Rhythms tied to biomechanical, articulatory or physiological set points varied least: walking cadence, syllabic rate, heart rate and individual alpha frequency (CV ≤ [0.16]).

### Statistical relationship between BBB rhythms

We statistically tested the hypothesis that BBB rhythms form a hierarchy with a constant ratio between neighboring rhythms (Fig. 2B, bottom). In other words, we examined whether rhythms across modalities followed an arithmetic progression on a natural-logarithmic scale.

We tested two main hypotheses: Buzsáki ^2,19^ proposed a ratio of about *e*: 2.72; as an alternative, Klimesch ^1^ alternatively proposed the golden rule (φ: 1.618) to define the bandwidth between adjacent rhythm classes. The *e* and golden rule would produce rates at which rhythms can be maximally decoupled, thus avoiding interference between independent processes. Next to these, we included three additional models: one defined by harmonic (integer ratios) relationships across modalities; a common, empirical ratio, estimated from all other participants (leave-one-participant-out); and a tailored model (‘free’) capturing participants’ own ratio.

While the *e* and harmonic ratio fitted poorly (RMSE = .62 and 1.24 log units (median across participants; Fig. 2B bottom), the φ ladder (RMSE = .43; *p* = .038) and the common empirical ratio (RMSE = .44; *p = .041*) fitted equally well (ΔBIC between these two best models = 4.3). Group BIC strongly disfavored individual ratios (190.4), ratio 2 (212.4) and ratio e (592.6).

### BBB rhythm classes

As some rhythm modalities clustered in rather close proximity to each other (e.g., cardiac, tapping and walking rates: 1.4-1.6Hz), we repeated the comparison creating rhythm ‘classes’ (Fig. 3). When the same data were summarized as four rhythm classes (pupil and breathing; tapping, heart, saccades and walking; speech; alpha), the best common step became 3.39 (Fig. 3A; RMSE = .21). While the fit to the *e* ladder model was modest (RMSE = .33; *p* = .035), the ΔBIC favored the ‘fitted’ model (ΔBIC = 97).

**Figure 3.**
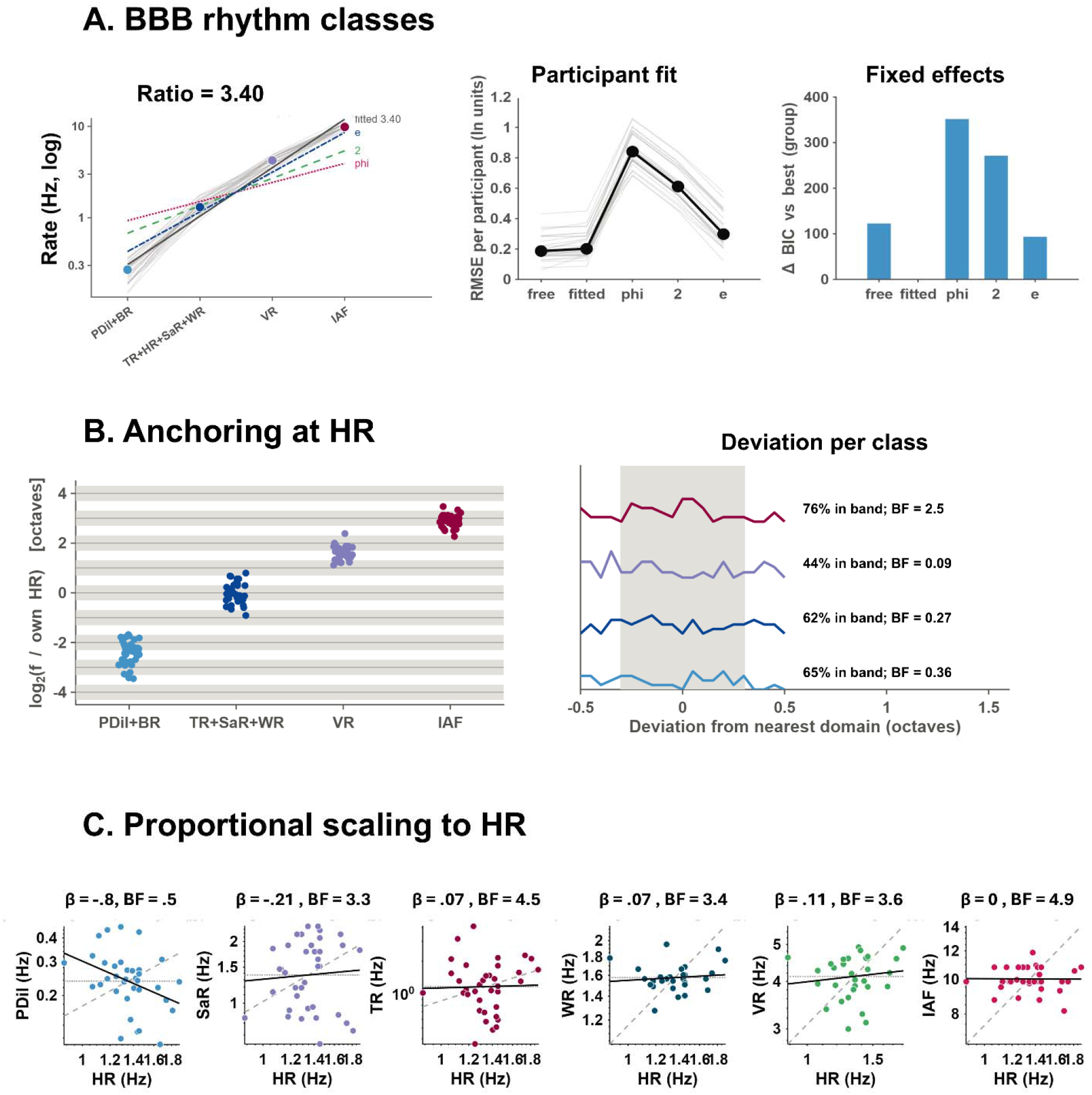
Rhythm classes in BBB rhythms. A, left: Rates of rhythms grouped per classes (<.5Hz difference), displayed on a logarithmic axis and ordered by group median. We calculated the log-ratio between adjacent rhythm classes and compared five descriptions of each participant’s rates across modalities: an individual model (‘free’); a common empirical model (leave-one-participant-out; ‘fitted’); three fixed ladders with ratios set to φ (phi: 1.618), 2 (harmonic) and ‘e’ (2.41). Each grey line represents one participant. Colored dots display the group mean per modality. black line: average constant-ratio fit (ratio per step 3.40). A, middle: root-mean-square deviation of each participant’s log-rates from five ladders: individual ratio (free), common ratio estimated from the other participants (fitted), and fixed ratios φ, 2 and e. Grey: participants; black: medians. A, right: group evidence for each ladder: summed BIC relative to the best model (lower is better; differences > 10 are strong evidence). B: Position of each rhythm class in octaves relative to the participant’s own heart rate. Grey bands: domains of Klimesch’s binary hierarchy (heart rate × 2ⁿ) with their bandwidths (±0.31 octaves). Heart rate is excluded from the class containing it. G: Distribution of deviations from the nearest domain per class. Grey area: within band (61% expected by chance). Labels: proportion in band and one-sided Bayes factor for a proportion above chance (BF+0). C: proportional scaling to HR. Y-axis: each modality rhythm in Hz; X: HR in Hz; both on log axes, one dot per participant. The solid black line is the fitted power law; the dashed gray line shows what β = 1 would look like (ratio conserved), and the dotted horizontal line shows β = 0 (independence). Both are drawn through the group means.

### Scaling law to individual HRs

Next, we tested Klimesch’s prediction according to which each person’s heart rate sets the scale for defining other rhythm classes. In this view, rhythm classes lie at HR multiplied or divided by powers of two.

In a first approach, we defined frequency bands spanning from the domain centre multiplied by φ/2 to the centre multiplied by 2/φ, i.e. about ±0.31 octaves. Relative to each participant’s own HR, the median class positions were −2.36 octaves for pupil and breathing, −0.06 for tapping, saccades and walking, +1.62 for speech and +2.93 for alpha (Fig. 3B). Klimesch’s binary hierarchy predicts whole numbers: −2 (a quarter of heart rate), 0, +2 and +3 (eight times heart rate). The proportion of participants inside the predicted bands was at chance for pupil and breathing (65% vs 61%; Bayes Factor+0 (BF) = 0.36) and for the motor class (62%; BF+0 = 0.27), and below chance for speech (44%; BF+0 = 0.09). Only alpha exceeded chance, and only anecdotally (76%; BF+0 = 2.5). Its ratio to HR (median 7.61) was close to the predicted 8 (BF10 = 2.8 for a deviation from 8; inconclusive evidence).

Finally, we tested the hypothesis of proportionality between rhythms in each modality and individual HRs. None of the rates scaled with HR (Fig. 3C). Most exponents were close to zero: walking (β = 0.07, 95% CI [−0.16, 0.35]), speech (β = 0.11 [−0.15, 0.45]), tapping (β = 0.07 [−1.22, 1.07]), and individual alpha frequency (β = −0.00 [−0.20, 0.14]), while the BF01 was >3 for most rhythms, thus rejecting the hypothesis of proportional scaling. Pupil rate was the only modality showing a negative exponent (β = −0.80 [−1.47, −0.28]), although the corresponding evidence was inconclusive (BF01 = 0.5, i.e. BF10 = 2) and did not survive correction for seven comparisons.

To exclude the possibility that simpler linear associations between raw (not log) modality rates would describe the relationship across BBB rates, we further tested whether individual BBB rates are linearly correlated with individual HR. While the power law described above asks *by what exponent* walking rate scales with HR (WR = c·HR^β), a pearson correlation on raw rates asks whether walking rate rises linearly with HR (WR = a + b·HR). Pearson’s linear correlations showed no significant associations between individual BBB rhythms and HRs at rest (Suppl. Fig. 6C), excluding the possibility that the observed rates linearly scale to one central modality. Results support prior observations ^42^.

### Exploratory analyses – Part I

We explored the hypothesis that interoception modulates the relationship between BBB rhythms. If so, one would identify at least two groups: high- and low-interoception clusters, characterized by stronger or weaker BBB inter-dependency.

The rationale was the following: if BBB rhythms are inter-dependent, their relationship should be consistently quantifiable. In turn, the statistical dependence across modalities should enable predictions: despite inter-individual variation in BBB rhythms, one should be able to predict individual rhythms in one modality leveraging inter-individual variation in the remaining modalities.

### Cross-modal rate deviation

We constructed a log-ratio representation for each participant, quantifying how consistently each participant’s physiological, neural, and behavioral rates co-vary relative to the group. The log-ratio removes the absolute scale differences between modalities but allows capturing the relative structure of each individual’s rate configuration. To this end, we computed pair-wise log ratios across participants and modalities and obtained a ‘deviation score’ (root-mean-square difference between each participant’s log-ratio profile and the group median profile) to quantify the distance between each participant’s log-ratio profile and the group median profile. In turn, small deviation scores indicate that a participant’s cross-modal rate structure conforms to the group-typical pattern; instead, large values indicate deviation from the group pattern and identify a more idiosyncratic profile.

To interpret these individual scores, we assessed the association between cross-modal rate deviation scores and individual MAIA scores via Spearman rank correlation (Fig. 4A and 4B left panel). We hypothesized that individuals with high interoceptive awareness (e.g., ‘Noticing’ and ‘Body listening’ subscales) would show more consistent BBB rate profiles.

**Figure 4.**
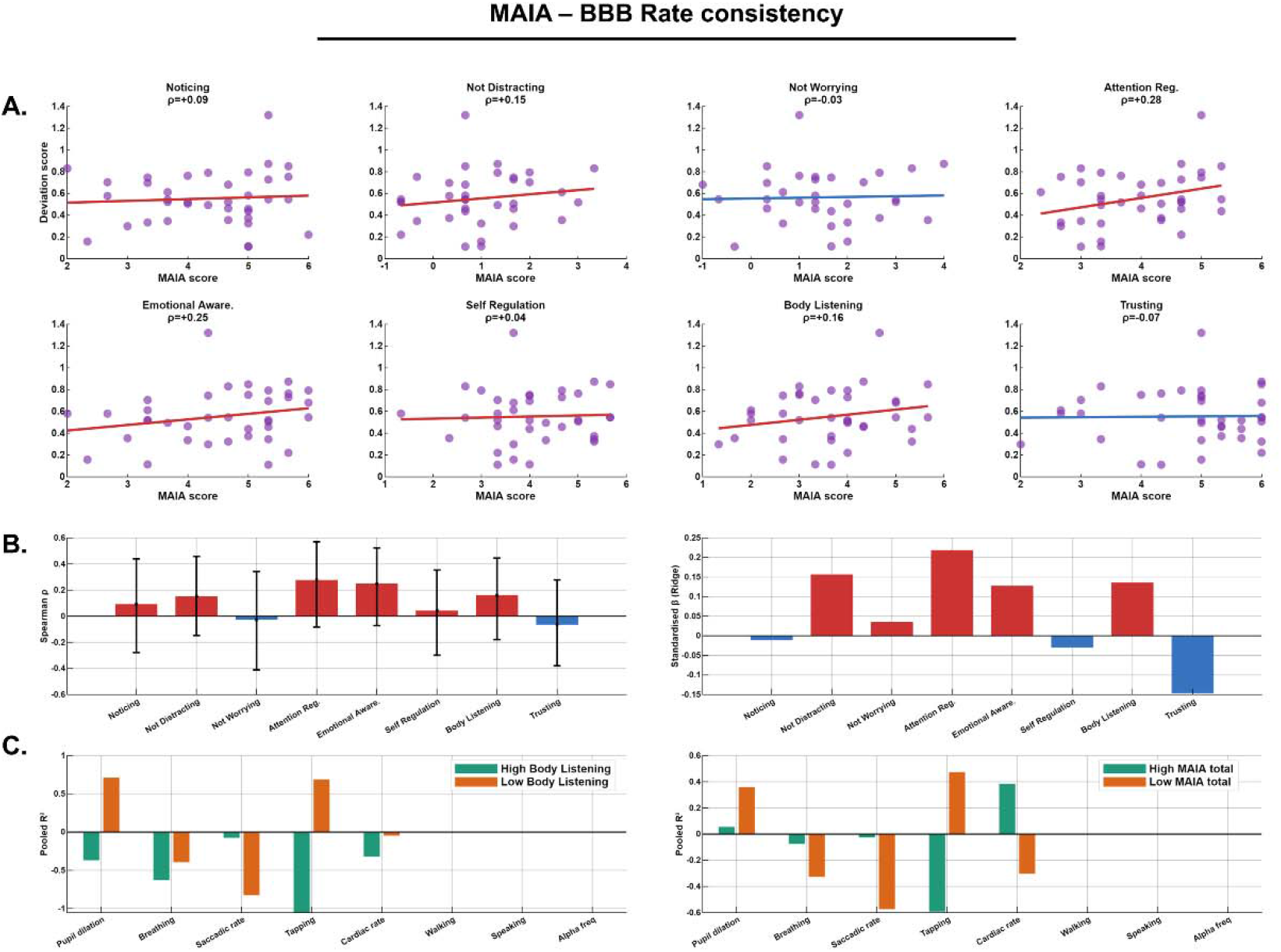
Cross-modal rate consistency and MAIA. A: Spearman correlations between cross-modal rate deviation scores and individual MAIA sub-scale. Each dot represents an individual. The colored line represents a positive (red) or negative (blue) association. B: represents associations drawn in A at the group level, adding Bootstrap 95% confidence intervals (black intervals). C: ridge regression model predicting deviation score from all eight MAIA subscales simultaneously. In red, positive associations; in blue negative associations. C-D: cross-modal prediction accuracy in high-(green) and low-(orange) Body-listening (left) and MAIA total score (right). Plots represent the pooled R2 across individuals within the same group.

Contrary to our a priori hypothesis, Spearman correlations between MAIA subscale scores and the cross-modal rate deviation score were predominantly positive, suggesting that higher interoceptive awareness was associated with *greater* deviation from the group-typical ratio profile — that is, more idiosyncratic cross-modal rate configurations — rather than more consistent ones (Fig. 4A and B (left)). The strongest positive correlations were observed for Attention Regulation (ρ = +0.28) and Emotional Awareness (ρ = +0.25), followed by Not Distracting (ρ = +0.15), Body Listening (ρ = +0.16), and Noticing (ρ = +0 09). Self-Regulation showed a near-zero association (ρ = +0.04). The only subscales exhibiting correlations in the hypothesized negative direction were Not Worrying (ρ = −0.03) and Trusting (ρ = −0.07). After Benjamini–Hochberg FDR correction for multiple comparisons across the eight subscales, no correlation reached statistical significance; a result we suggest to interpret in the context of limited statistical power available at the present sample size (rather than as unambiguous evidence of absence). Bootstrap 95% confidence intervals, displayed in Fig. 4B (left), were wide for most subscales and spanned zero, confirming that individual associations might not be reliably estimated at *N* = 37. Nonetheless, the directional consistency of positive correlations across six of the eight subscales — with no subscale showing a meaningful negative correlation — may suggest a modest pattern.

### Unique Contributions of MAIA Subscales to cross-modal rate deviation

Next, we employed a ridge regression model to identify which MAIA subscales uniquely and independently predicted deviation score after accounting for inter-subscale correlations.

The Ridge regression model predicting deviation scores from all eight MAIA subscales simultaneously (Fig. 4B, right panel) broadly reproduced the pattern observed for the bivariate correlations whilst additionally revealing the independent contribution of each subscale after partialing out inter-subscale shared variance. Attention Regulation showed the largest positive standardized coefficient (β ≈ +0.22), indicating that greater capacity to regulate attentional focus toward bodily signals was uniquely associated with more idiosyncratic cross-modal rate profiles. Not Worrying (β ≈ +0.15) and Emotional Awareness (β ≈ +0.13) also showed positive unique contributions. Self-Regulation and Body Listening contributed smaller positive coefficients.

Trusting emerged as the only subscale with a clearly negative standardized coefficient (β ≈ −0.13), indicating that, when accounting for shared variance with the other subscales, greater trust in bodily sensations was uniquely associated with *more* consistent cross-modal rate profiles — the only MAIA dimension that aligned with the original hypothesis. The leave-one-out cross-validated performance of this regression model was generally low (LOO-R2 <.1), consistent with the weak bivariate associations, indicating that the reported coefficients should be treated as descriptive rather than confirmatory.

### Cross-Modal Prediction Accuracy in High-versus Low-Interoception Groups

Finally, we tested whether interoceptive awareness moderated cross-modal rate predictability. To this end, participants were divided in two groups via a median split on Body Listening. Next, we trained a model to predict one modality from inter-individual variation in the other modalities (20% test set). Within each subgroup, a leave-one-modality-out prediction was performed. Prediction accuracy was evaluated via Monte Carlo random subsampling cross validation with 1000 iterations, and we pooled per-fold predictions to compute an R² per subgroup.

Stratification by Body Listening (Fig. 4C, left panel) revealed a clear dissociation in prediction accuracy between groups. Participants in the low-Body-Listening group showed higher — in some modalities markedly positive — pooled R² relative to those in the high-Body-Listening group. The most pronounced difference was observed for pupil dilation rate, where the low-Body-Listening group yielded a pooled R² of approximately +0.65, compared with approximately −0.20 in the high-Body-Listening group. For tapping rate, a similar pattern emerged, with the low-Body-Listening group showing R² of approximately +0.60 against deeply negative values (approximately −0.85) in the high-Body-Listening group. Remaining modalities — breathing rate, saccadic rate, cardiac rate, walking rate, speaking rate, and alpha frequency — showed more modest or near-zero R² in both groups, consistent with the lower inter-individual variability of those modalities.

We repeated the same procedure by stratifying the group according to the MAIA total score (Fig. 4C, right panel) and observed an overall comparable pattern. The low-MAIA-total group showed positive R² for pupil dilation (R² ≈ +0.40), tapping rate (R² ≈ +0.40), and, to a lesser extent, saccadic rate and speaking rate, whereas the high-MAIA-total group showed near-zero or negative R² for most targets, with cardiac rate representing a partial exception (R² ≈ +0.40 in the high-MAIA group).

We acknowledge that with a within-group sample size of approximately 18 participants, per-modality R² estimates are susceptible to instability and should therefore be interpreted with appropriate caution. Nonetheless, the consistency of the directional pattern across both grouping criteria (Body Listening and MAIA total) and across more than one modality strengthens the interpretation that the effect might not be exclusively attributed to sampling variability.

### Exploratory analyses – Part II

We further explored the link between individual behavioral rates (walking and tapping) and HRs. To this end, we adopted two complementary approaches: cross-correlations and phase analyses. Inspection of the time-courses of tapping, walking, and HRs (Suppl. Fig. 6A-B) suggest participants display at least two behaviors: while some individuals tend to tap and walk in close proximity to their HRs (e.g., if one’s HR is 1.4Hz, one walks at a rate of 1.2 – 1.6Hz; Suppl. Fig. 6A), others show rather independent rates (Suppl. Fig. 6B). To statistically quantify time-varying inter-dependency between behavioral and HRs over time, we performed cross-correlation analyses on 9 lags (−4,0,+4) utilizing an alpha level of .05. We observed large intra-individual variability in *rho* values (Suppl. Fig. 7), suggesting that the relationship between behavioral rates and HR time-courses fluctuated widely during the recording. Next, we calculated the proportion of significant correlations (*p* < .05; irrespective of their sign (positive / negative *rho*)) with respect to the entire recording length, per participant. Correlation frequency fluctuated in a range between ∼10-37% in TR across participants (Suppl. Fig. 7A), yielding a median correlation frequency of ∼18%. The correlation frequency in WR was slightly lower, fluctuating between ∼9-28% across participants, and yielding a median correlation frequency of ∼16% (Suppl. Fig. 7B). These observations further strengthen the notion that there are rather sparse linear associations between individual behavioral and HRs ^42^.

Next, we tested whether the cardiac phase would modulate movement onset timing. To do so, we identified R-peaks in ECG (see 2.4.4) and discretized the time-windows between successive R-peaks defining the cardiac cycle (0 - 360°; Suppl. Fig. 6D, top panel). Finally, we mapped tapping and step onset timing to the cardiac cycle across the recording, obtaining a distribution of movement onset timing per phase-bin (Suppl. Fig. 6D, bottom panels). To statistically assess whether movement onsets were more likely to occur in a specific phase of the cardiac cycle we employed circular statistics (circular statistics toolbox) and calculated the mean vector length per participant and group. Despite some individual preferences for clustering tapping or step onsets to a specific phase of the cardiac cycle (see single participant data in Suppl. Fig. 6D), no consistent pattern was observed at the group level (*p >* .05) suggesting no linear relationship between movement onset timing and cardiac phase.

## Discussion

In a recent perspective article, Buzsáki ^2^ hypothesized the existence of a *dynamic* cross-scale frequency architecture encompassing biological periodicities in the body and the brain. Embracing evidence that logarithmic properties are found in brain structural and functional parameters (from the size of dendritic boutons to synaptic weights and neuronal firing) ^22,43^, as well as in space, time and memory processing (Weber-Fechner law ^16–18)^, Buzsáki ^2,19^ proposed the body-brain frequency architecture to display an arithmetic progression on a natural logarithmic scale. In this view, the center frequencies and bandwidths of successive rhythm classes (from ultra-slow to fast) would be described by a ratio of about *e*: 2.72.

In a similar perspective, Klimesch ^1^ hypothesized body-brain-behavioral rhythms to progress along a ladder whose bandwidths between adjacent rhythms is defined by the so-called ‘golden mean’: φ = 1.618. In both perspectives, body-brain-behavioral (BBB) rhythms are not random; they form a frequency architecture where rhythm classes (Buzsáki) and individual rates (Klimesch) are defined by irrational ratios with other periodicities, thus ensuring harmonic interference to be minimized in function of communication efficiency. While avoiding ubiquitous phase coupling across modalities, cross-frequency calibration mechanisms would adaptively modulate phase-amplitude interactions across rates and the brain with the scope to optimize information sampling and action coordination. That is, while each rate is independent and maximally de-coupled from others, modalities can dynamically engage in cross-frequency interactions.

### A frequency architecture in body-brain rhythms

Here, we directly addressed some of the main implications arising from these perspectives by recording multimodal body-brain-behavioral data at rest, in action and perception. With a unique combination of smart wearable technologies ^44^, we were able to characterize individual endogenous rhythms in pupil dilation, breathing, cardiac, and neural activity, as well as in spontaneous behaviors (from tapping to walking and speaking). Next, we tested whether rhythms across the body, brain and behavior are randomly distributed in frequency or organized according to a scaling law, as proposed by recent frameworks of body-brain timing. Three observations argue for a non-random organization. First, each rhythm occupied a characteristic frequency, from about 0.25 Hz (pupil, breathing) to about 10 Hz (alpha). Inter-individual differences were lognormally distributed for the most variable rhythms, and inter-individual variability grew sub-linearly with mean frequency (slope = 0.57). This is consistent with multiplicative generative processes and with the proposal that the variability of brain–body rhythms increases with their mean frequency ^45^. Second, rhythms grouped into four classes: slow autonomic (pupil, breathing, about 0.25 Hz), cardio-motor (heart, tapping, walking, saccades, about 1.4–1.6 Hz), syllabic (speech, about 4 Hz) and alpha (about 10 Hz). The center frequencies of these classes advanced by an approximately constant ratio (3.4), a progression that was best captured by *e*; in line with the natural-logarithmic hierarchy of brain oscillations ^19^ and its extension to body-brain rhythms ^2^. Third, at the finer level of individual modalities, successive rhythms were separated by a ratio of 1.63; indistinguishable from the golden mean. Klimesch and colleagues proposed exactly this spacing as optimal: an irrational ratio minimizes spurious phase synchronization between neighboring frequencies while still allowing transient functional coupling ^1,36,37^. The two frameworks thus appear complementary, each describing a different level of one architecture: e-spaced classes partition the frequency axis into broad timescale domains, and golden-mean spacing governs how neighboring rhythms are arranged. Altogether, results confirm the existence of a cross-modal frequency architecture linking biological periodicities in the body and the brain with behavioral rhythms ^1,2,38,46^.

### Behavioral rhythms extend the body–brain hierarchy

Spontaneous behaviors did not occupy arbitrary frequencies. Self-paced tapping, walking and saccades during free viewing fell within the same class as the heartbeat, and syllabic speech rate fell within the next class. Self-generated movement therefore adopts frequency positions within the body–brain hierarchy rather than outside it, suggesting that the architecture extends from physiology to action. This extension, however, did not take the form of rigid coupling within individuals. Behavioral and neural rates did not scale with an individual’s own heart rate (exponents β ≈ 0; BF01 = 3.3–4.9), and individual rhythms did not sit at fixed octave positions relative to the individual’s heart rate (BF ≤ 0.36 for three of four classes; anecdotal evidence only for alpha, BF = 2.5). These results, along with the absence of linear correlations between individual behavioral (tapping and walking) rates and HRs, and of consistent phase clustering of behavior (taps and steps onsets) with the cardiac cycle (see for large reviews ^33,34^), replicate and extend earlier null associations among spontaneous motor tempo, heart rate and alpha frequency ^42^. These observations may indicate that the architecture constrains *which timescale domain* a rhythm occupies, not its exact value in a given person. Within each domain, individuals vary independently, and this variability depends on how constrained the rhythm is. Walking cadence, syllabic rate, heart rate and alpha frequency were tightly distributed, whereas self-paced tapping and the slow autonomic rhythms varied most.

Ideally, muscle relaxation during systole would optimize blood flow into the skeletal muscle and reduce cardiac afterload – a phenomenon known as cardiometabolic efficiency ^47^. While individuals can learn to time foot strikes to occur during cardiac diastole ^48,49^, there are large stride-to-stride fluctuations in gait, which exhibit long-range fractal (1/f) correlations ^50^ and may consequently fail to cluster at a precise phase of the cardiac cycle.

Altogether, null results on the binary link between cardiac and behavioral rates suggest that these measures may reflect partially independent components of the body–brain system, rather than a single, unified *pacemaker*. But then, how does the body–brain system generate and maintain an internal sense of time to guide perception and action?

### Implications for an internal representation of time

Traditional psychological models ^13,17,26–28^ propose the internal sense of time to rely on an ‘observer’ (or *accumulator,* or *perceptual store* ^28^) integrating the number of pulses emitted by a stable internal *pacemaker*. Embodied models proposed a strong role of the heart in modulating temporal processing, but produced mixed results ^35,42,51–54^, ultimately resonating with other evidence assessing time judgements in relation to various physiological parameters (e.g., not only cardiac, but also respiratory rates and body temperature ^55,56^). These views are somehow extended by neural oscillation models ^16,29–32^, grounding the ‘pacemaker’ perspective into neurobiological mechanisms: not one, but an array of pacemakers, resembled by endogenous neural oscillations across multiple timescales, were thus proposed to track the passage of time, accounting for the rich set of timescales we experience.

Our data argue against a single body-based pacemaker: intra- and inter-individual variability in heart rates did not propagate to other rhythms, as they would if one there anchor paces the system. The results are more compatible with accounts in which duration is read out from the relations among oscillators at multiple timescales ^16,29–32^. In fact, a log-spaced, golden-mean-separated set of rhythms may provide such a multi-scale reference, thus encompassing the wide range of experienced time. We propose that the body-brain frequency architecture may act as a temporal reference frame, scaffolding the coordination of self-paced movements, and against which elapsed time can be estimated. Because the frame is defined by the relative positions of rhythm classes, the temporal reference remains stable across individuals while tolerating considerable individual variation in each component.

As we did not assess time perception per se, this remains a hypothesis that generates testable predictions. First, individual subjective tempo and duration judgements should be influenced by individual’s rhythm classes. Second, manipulations that shift whole classes, such as arousal, pharmacological or metabolic challenges known to distort subjective time (e.g., ^28^), should distort time estimates in proportion to that shift.

Traditional perspectives on time cognition partially neglected early evidence documenting that emotional states, attention, stress ^28^ and dietary ^57^ choices were associated with subjective distortions of the internal sense of time in a comparable way to drug manipulations: while methamphetamine increased the *internal pacemaker* speed (induced a leftward shift in the psychometric function for duration estimation) ^58^, carbohydrate intake slowed down the pacemaker (leading to a rightward shift of the psychometric function) ^57^. These results suggest that complex mind-body states may influence the perception and estimation of time ^15,28^. In turn, multiplexed, cross-frequency inputs across the body ^59^ and the brain ^60^ may communicate *change,* ultimately allowing for an emergent sense of time. In this view, not a single master clock, but distributed clocks at different frequencies should be integrated. This perspective adds an important (modulatory?) component to *accumulator* (or ‘observer’) regions traditionally ascribed to timing processing networks in cortico-subcortical ^16,61–63:^ interoceptive networks sensing changes from the viscera ^34,64–66^. Among these, the almost ubiquitous activation of supplementary motor area and insular cortex in timing tasks ^67,68^ , along with the involvement in processing ascending body signals ^24,52,68^, may suggest a strong involvement in the integration of cross-modal sensory input and in the subjective feeling of time.

### From architecture to dynamics

We discussed that the existence of hierarchical scaling laws governing the endogenous rate of biological and behavioral signals may expand traditional psychological ^13,17,26–28^ and neural ^16,29–32^ models of time processing, enriching them with an *embodied* perspective that links the repertoire of *subjective* time with dynamic body-brain-behavioral states ^2^.

The empirical demonstrations presented here – though in a small sample of young, healthy participants – suggests that a scaling law may coordinate the body-brain system: a complex, multi-scale oscillatory system whose cross-frequency interactions may be calibrated to optimize information processing and motor control ^33,38–40,69^. Resonating with earlier hypotheses, we strengthen the view that the body-brain frequency architecture may ultimately provide a temporal reference frame – a scaffold to adaptively coordinate perception and action via an emergent *sense* of time ^33,38^. In this view, the multi-scale organization may – on the one hand – provide a mechanism to ensure homeostatic stability; and - on the other hand – enable behavioral flexibility: a dynamic equilibrium ^70,71^ maintained by cross-frequency modulatory interactions.

However, the architecture described here is a rather static phenomenon: we characterized *where* rhythms sit in frequency; not how they interact moment to moment with other biological and behavioral rhythms. In order to test whether an embodied temporal reference frame is actually employed to coordinate movement requires showing that rhythms adjust to one another in real time; for instance, through transient phase synchronization dynamics within and across classes. Coupled-oscillator models and the estimation of coupling functions ^72,73^ may provide the right tools to test this in continuous multimodal recordings in future work.

### A link between interoception and BBB rhythms?

A modest degree of inter-individual variability in interoceptive dimensions (as measured by the MAIA-2 scale ^74^) allowed asking the question of whether interoception modulates the body-brain frequency architecture.

Interoception is thought of as the ability to process, regulate, predict, and integrate visceral sensations with contextual information from the external sensory environment ^23,66,75–79^. Interoception is increasingly recognized as a fundamental modulator of neurocognitive functions and mental health ^33,41,65,80–82^. In a series of exploratory analyses, we asked (i) does inter-individual variation on sub-dimensions of interoception modulate individual cross-modal rate deviation (i.e., a score quantifying the distance between each participant’s cross-modal rate profile and the group median)? Secondly, (ii) do MAIA subscales uniquely and independently predict deviation scores after accounting for inter-subscale correlations? Finally, (iii) can we predict individual body-brain-behavioral rates from inter-individual variation in other modalities?

Taken together, the current results revealed a dissociation between interoceptive awareness and cross-modal rate predictability: while we hypothesized that individuals with high interoceptive awareness (e.g., ‘Noticing’ and ‘Body listening’ subscales) would show more consistent BBB rate profiles, results suggested trends in the opposite direction, whereby higher interoceptive awareness was associated with more idiosyncratic cross-modal rate configurations. Hence, individuals with lower interoceptive awareness exhibited more stereotyped, group-typical cross-modal rate profiles that are better captured by a population-level prediction model. Trusting (i.e., trust in bodily sensations) was the only subscale uniquely associated with *more* consistent cross-modal rate profiles. However, given the low statistical power at the present sample size, the median splitting of the group, and the high number of missing data, most observations should be interpreted as descriptive rather than conclusive and motivate future research with larger samples.

### Limitations and future directions

In this study, we aimed at characterizing endogenous rhythms in the body, brain and behavior and assessing their statistical relationship. We strongly encourage future research to try and replicate our study, and further address the research questions emerging here in larger samples and with complementary methodologies (e.g., MEG). Here, the choice of employing wearable devices has increased the overall ecological validity of the paradigm; at the same time, however, it has limited the range of possible computational approaches to better characterize body-brain-behavioral dynamics. In particular, the low electrode density of the EEG system, along with the poorer signal-to-noise as compared to standard EEG recorded in magnetically-shielded rooms, hindered not only spatial localization of observed electrical fluctuations on the scalp but prevented modelling complex interactions in the body-brain system. We encourage future research to embrace the high-dimensionality offered by body-brain investigations and characterize how dynamic body-brain interactions modulate cognition ^33,38^. In doing so, future studies would enrich current research – which most exclusively focused on rest and spontaneous behaviors, neglecting information processing in perceptual tasks. For instance, recent conceptualizations ^39,40^ suggest discretizing body-brain interactions into low-dimensional state spaces whose trajectories span micro-, meso- and macro-level dynamics (from subseconds, to minutes to days and years). Such dynamical system approach promises to better capture the inherent complexity of nested interactions across temporal scale, advancing our understanding of how they coordinate information processing, perception and action from healthy to pathological conditions ^39,40^.

## Conclusions

Combining multimodal body-brain-behavioral data recorded from a unique set of wearable devices at rest and in perception and action, we characterize a cross-scale frequency architecture linking biological periodicities with behavioral rhythms. The architecture displays an arithmetic progression on a natural logarithmic scale, extending such property from the brain and other biological system to a body-brain dynamic system.

## Supplementary figures

**Suppl. Fig. 1.**
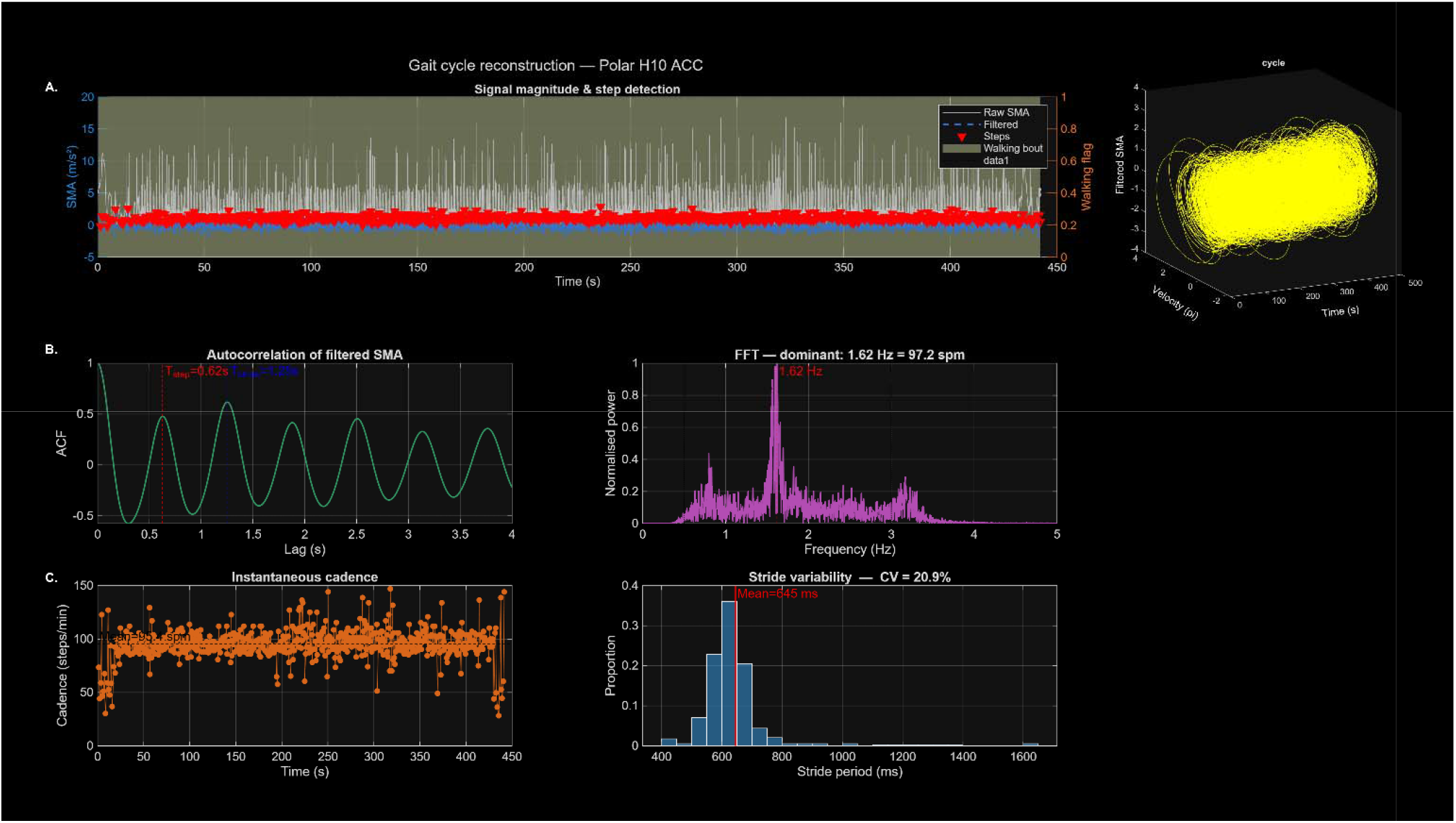
Gait cycle reconstruction A: signal magnitude area (SMA) expressed in m/s2 (y-axis) plotted over time for the raw SMA signal and the frequency-filtered SMA (dashed blue line). Red triangles indicate step onset times and gray shaded areas mark the time in which walking was detected. On the right, the gait cycle projected on a 3D-axis: time, velocity and amplitude. B: autocorrelation of filtered SMA (ACF; y-axis) plotted over time-lags (s; x-axis). On its right, the Fourier spectrum of the SMA signal (frequency (x-axis) by normalized power (y-axis)). C: Instantaneous cadence (y-axis: steps per minute) over time (in s; x-axis) and the mean steps per minute (dotted black line). On its right, the stride variability displayed as the proportion of stride periods (in ms). Red vertical line indicates the mean stride period.

**Suppl. Fig. 2.**
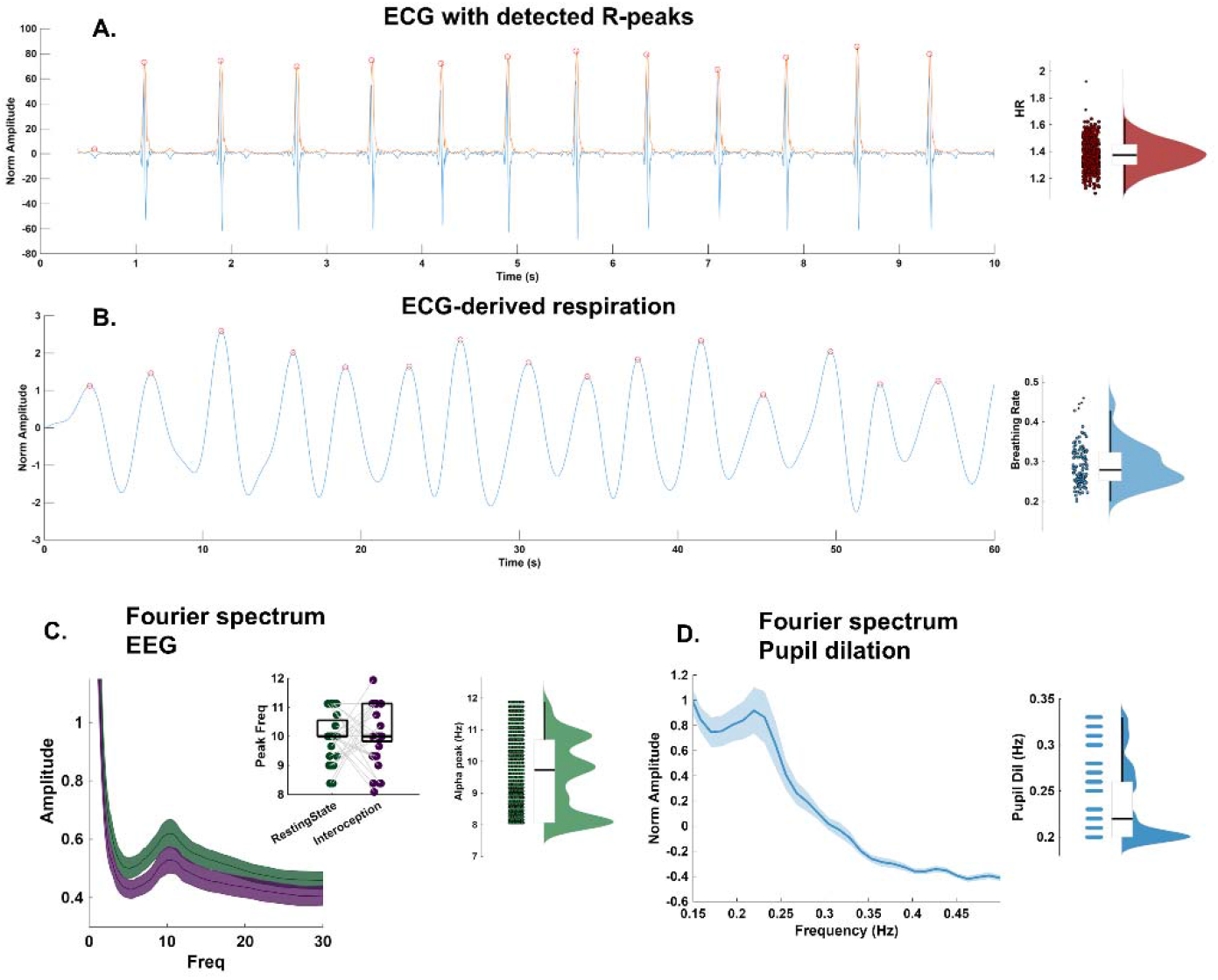
Example on body-brain rhythm identification. A: raw ECG signal (blue), its amplitude envelope (red) and R-peaks (red circles). On the right, the distribution of inter-beat intervals defining the median heart rate (HR; black horizontal line). B: ECG-derived respiratory signal (blue) and identified inspiration peaks (red circles). On the right, the distribution of inter-peaks intervals defining the median breathing rate (BR; black horizontal line). C-D: Fourier spectrum for EEG signals (left; green for resting state and purple for interoceptive attention) and pupil dilation (right). Insert panels, in order from left to right: identified peak frequencies in EEG, distribution of alpha peak frequencies and median alpha frequency (horizontal line), distribution of pupil dilation peak frequencies and its median (horizontal line).

**Suppl. Fig. 3.**
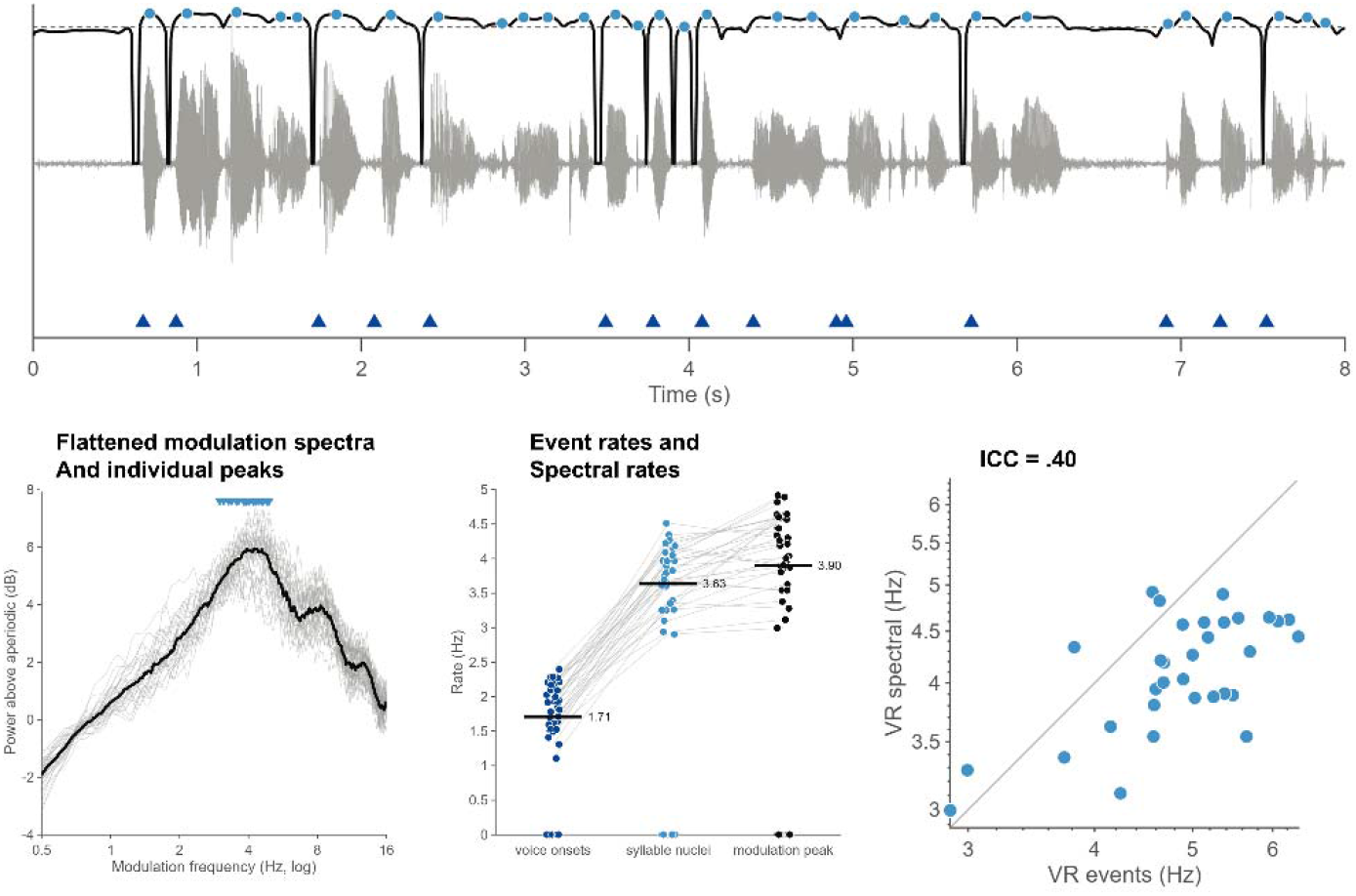
Speech rhythms. Top: 8-s excerpt of the waveform (grey), broadband intensity envelope (black) and detection threshold (dashed). Blue triangles: voice onsets; blue dots: syllable nuclei. Bottom left: modulation spectrum after removal of the aperiodic background. Grey lines for single-participants; black line for the group mean. Blue triangles show individual peaks. Middle: distribution of voice onset rates, syllable nuclei rates and modulation peak rates. Single dots and grey line per participant; horizontal black lines display the group mean. Right: inter-class correlation (ICC) between spectral and discrete event rates, in Hz.

**Suppl. Fig. 4.**
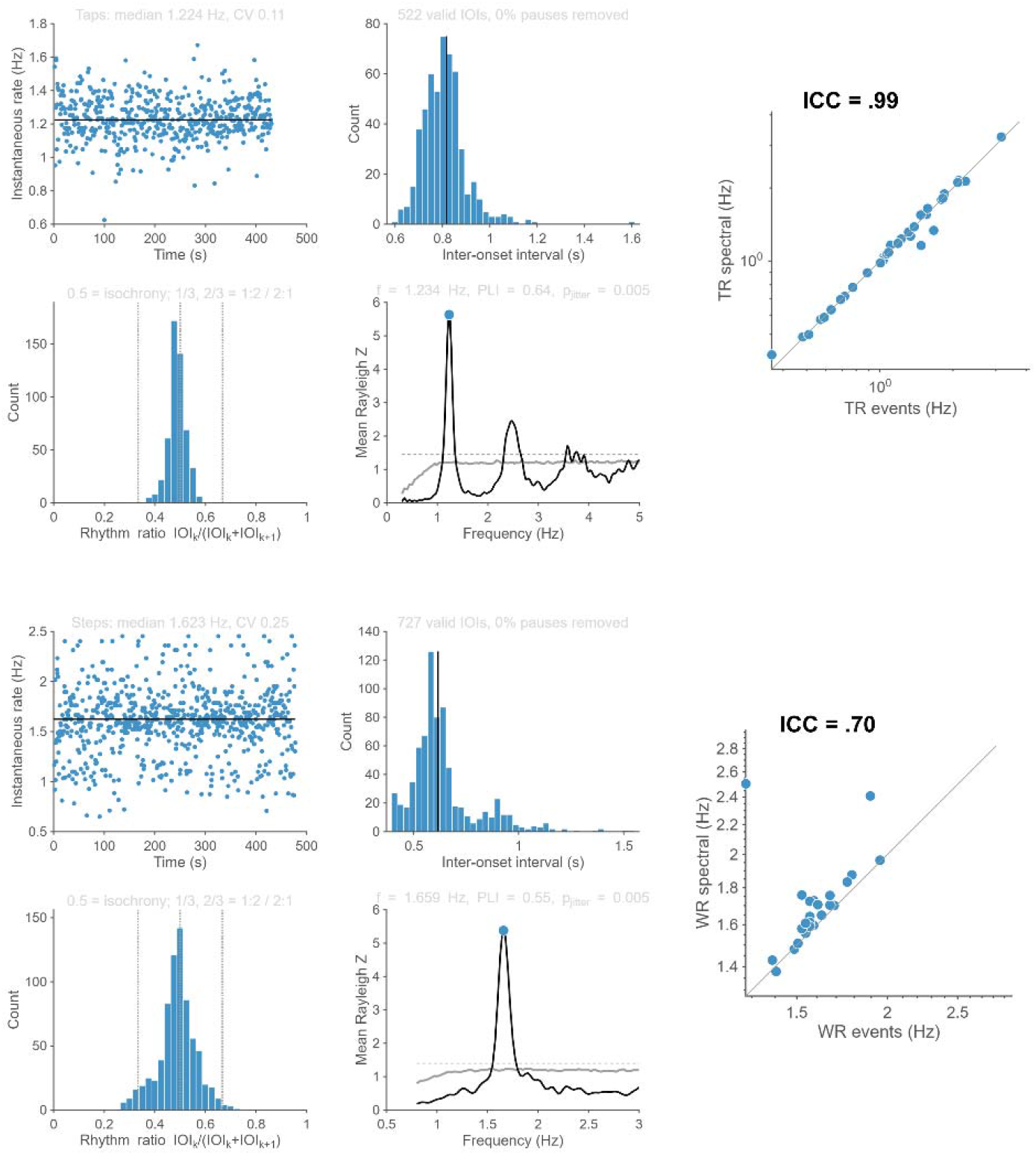
Rate and rhythmicity analyses for discrete tapping and walking signals. Top: tapping; Bottom: walking. Each have five panels: first, we computed the instantaneous frequency and the coefficient of variation from inter-onset interval time-series. Next, we assessed rhythmicity with a windowed Rayleigh test (frequency in Hz on the x-axis; and Rayleigh Z on the y-axis), tested against surrogates in which each event was displaced randomly by up to half the median interval. We inspected rhythm ratios between successive intervals as an additional, assumption-free signature of isochrony. Finally, we estimated the agreement between independent methods via intra-class correlation (ICC).

**Suppl. Fig. 5.**
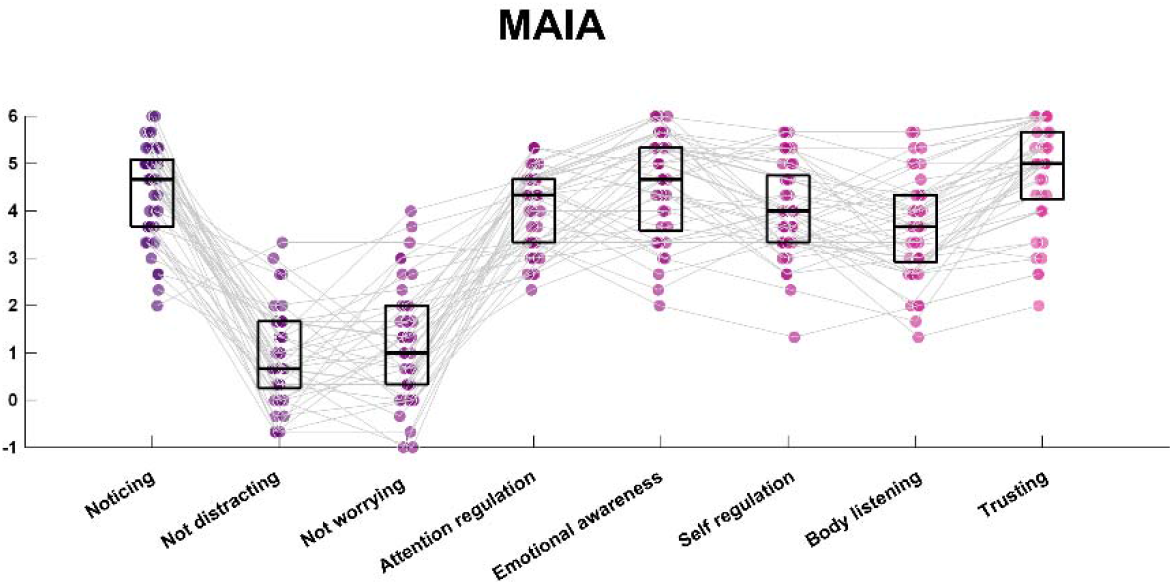
Individual MAIA scores. Individual Maia scores across subscales: noticing, not distracting, not worrying, attention regulation, emotional awareness, self-regulation, body listening, trusting. Horizontal lines connect individual scores across subscales.

**Suppl. Fig. 6.**
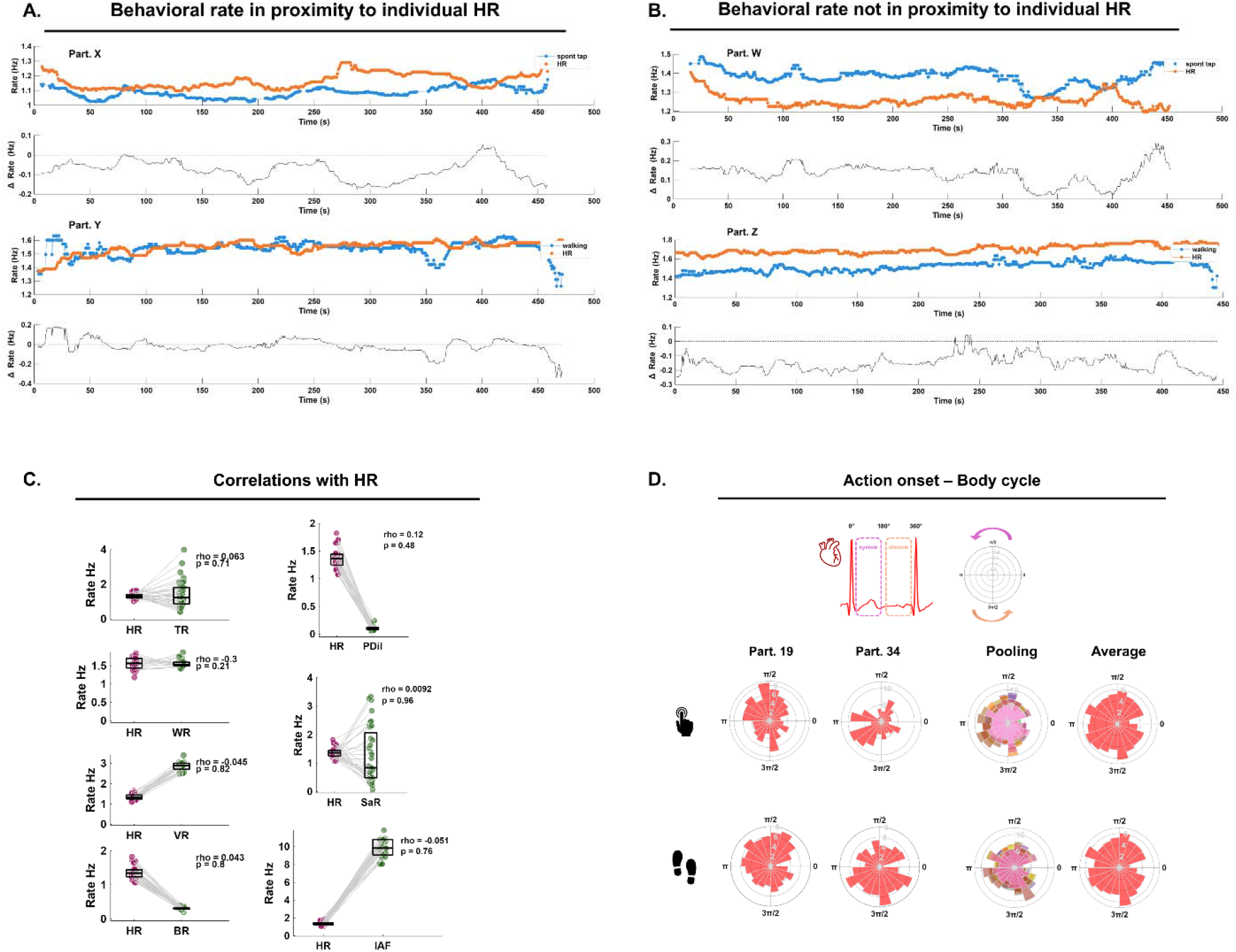
Linear relationship between BBB rates. A: examples of two participants (called here ‘X’ and ‘Y’) whose behavioral rate (tapping on top, walking at the bottom; blue dots) fluctuated in proximity to their heart rates (HR; red dots). The second and fourth panels from top to bottom show the difference rate between behavioral and HR. B: as in A, but now providing examples of two participants (called here ‘W’ and ‘Z’) whose behavioral rates did not fluctuate in proximity to their individual HRs. C: linear correlations between individual HR and body-brain-behavioral rates. In order from top to bottom, and from left to right: tapping rate (TR), pupil dilation rate (PDil), walking rate (WR), saccadic eye movement rate (SaR), voice onset rates (VR), breathing rates (BR) and individual alpha frequency (IAF). D: circular histograms providing the count of behavioral onsets to cardiac cycle: top for tapping and bottom for walking. From left to right: example data from participant N19, N34, pooling of data across participants (color-coded per participant), and group-level circular average.

**Suppl. Fig. 7.**
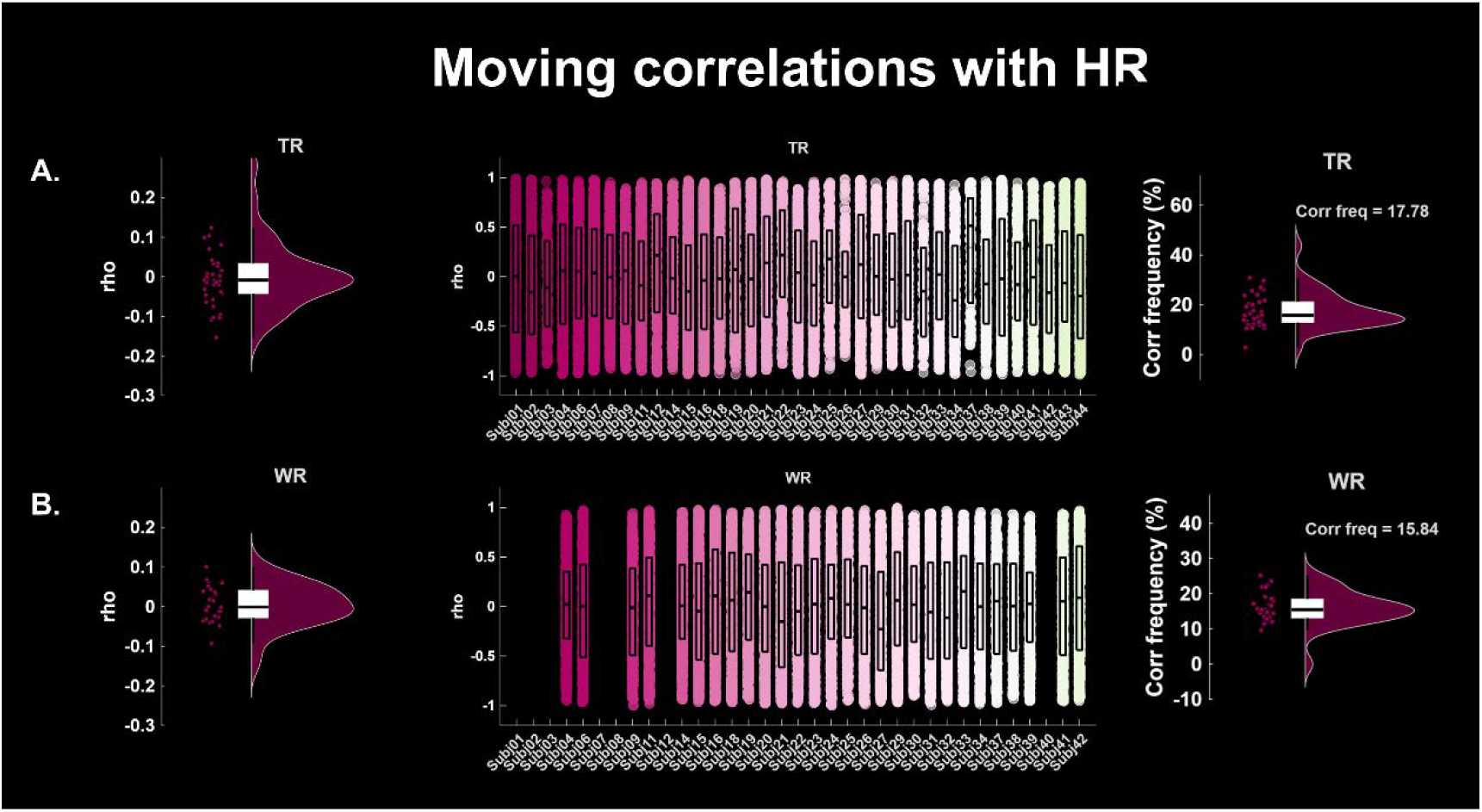
Cross-correlations between behavioral – heart rates. A: on the left, the distribution of mean cross-correlation values between individual tapping (TR) and heart rates (HR) at the group level (one dot per participant). In the middle, the distribution of cross-correlation values within each participant (x-axis).. On the right, the frequency (expressed as percentage) of significant cross-correlations at the group-level (one-dot per participant). Across panels, horizontal black lines report the median of the distribution. B: same as in A, but for the cross-correlation individual walking (WR) and HRs.

## Methods

### 2.1 Participants

Fifty participants took part in this pre-registered study ^44^ and signed written informed consent in accordance with the ethics committee of the University of Maastricht (OZL_255_97_07_2022) and the declaration of Helsinki. Technical difficulties reduced the original sample: recordings from one participant were not correctly saved; faulty synchronization across wearable devices led to loss of five datasets. A final sample of 44 participants was included in this study. The group comprised healthy young individuals recruited from the student population of Maastricht University (mean age = 24 years; age range: 19-37 years; 8 males). Of these, 13 participants had formal musical education (2-5 years). Participants reported no history of hearing deficits, nor psychiatric disorders. All participants received either monetary compensation or credit hours for taking part in the study.

### 2.2 Data Acquisition

Multimodal body-brain-behavioral signals were recorded simultaneously using a wearable sensor array (Fig. 1) and in a set of 10 tasks designed to maximize ecological validity.

All data streams were integrated via a custom-built application (described in §2.2.6) that assigned synchronized timestamps to each incoming signal, enabling post-hoc reconstruction of aligned time-series across modalities.

#### 2.2.1 Electroencephalography (EEG)

Continuous EEG was recorded using the MUSE S headband (InteraXon Inc., Toronto, Canada; https://choosemuse.com/), a dry-electrode device comprising four recording channels (TP9, AF7, AF8, TP10) referenced to one frontal electrode (Fpz). Despite its limited spatial resolution, MUSE 2 has been validated for use in research contexts ^83^. Raw EEG was streamed at 256 Hz via Bluetooth Low Energy (BLE) to the acquisition application. Electrode–scalp contact was verified prior to each session by visual inspection of signal quality indices provided by the MUSE software development toolkit (SDK).

#### 2.2.2 Cardiac Activity

Cardiac electrical activity was recorded using the Polar H10 chest strap (Polar Electro Oy, Kempele, Finland; https://www.polar.com/it/sensors/h10-heart-rate-sensor), which provides single-lead ECG at a sampling rate of 130 Hz. The Polar H10 has been validated against clinical-grade ECG devices and is widely used as a gold-standard wearable for heart rate variability (HRV) research ^84^. The device streamed data via BLE to the acquisition application.

#### 2.2.3 Respiration

Respiratory activity was estimated as the ECG-derived respiration (EDR) signal, leveraging the physiological dependence between respiration depth and rate and ECG fluctuations.

EDR has been widely utilized in biosignal processing (e.g., ^85^ ) and neuroimaging research ^86,87^. As our scope was not to map precise body-brain-behavioral coupling to the breathing cycle but to characterize intra- and inter-individual variabilities in body-brain-behavioral rhythms, the proposed method is thought to deliver a robust estimation of true respiratory frequency (see ^87^).

In short, the employed EDR algorithm involves the following steps: band-pass frequency filtering the ECG signal (5-30Hz with an IIR filter) and amplitude normalization to remove high-frequency noise and baseline wander, ultimately helping isolating the QRS complex; using a robust R-peak detection algorithm (Pan-Tompkins pipeline; ^88^); extract the R-wave amplitude at each R-peak location; reconstructing a continuous signal interpolating the R-wave amplitude using spline interpolation; band-pass filtering the signal in the respiration frequency band (.1-.5Hz with an IIR filter); mean-centering and flipping the polarity.

#### 2.2.4 Body Movement

Movement was continuously monitored via inertial measurement unit (IMU) embedded in the MUSE and POLAR bands. While the former provides triaxial acceleration and gyroscope data at 256Hz, POLAR provides acceleration at a sampling frequency of 50Hz. Body movement data were used for two main scopes: removing motion-related artifacts from EEG (see ‘EEG preprocessing pipeline’) and estimating individual walking rates via a gait reconstruction algorithm (see ‘Gait cycle reconstruction’).

#### 2.2.5 Eye movements and Pupil dilation

Binocular eye movements and pupil dilation signals were recorded using the Tobii Smart Pro 3 wearable glasses (https://www.tobii.com/products/eye-trackers/wearables/tobii-pro-glasses-3). The glasses recorded gaze position, pupil diameter, and blink events at a sampling rate of 100Hz. Tobii glasses have been widely used in experimental research across fields, including in psychology and neuroscience ^89,90^. The glasses were worn throughout the entire session and across tasks, and data was continuously recorded in Tobii’s native app. Prior to each session, a standard calibration procedure was performed to ensure gaze tracking accuracy. The calibration procedure was repeated if gaze tracking accuracy diminished during the experiment.

#### 2.2.6 Multimodal Data Integration and Synchronization

Data streams from MUSE and POLAR were integrated in real time using a custom acquisition application integrating the SDK of respective manufacturers. Each incoming data packet was tagged with a common network clock timestamp when received. Post-hoc, a dedicated preprocessing pipeline integrated and temporally aligned multivariate time-series by interpolating each stream onto a common 256Hz sampling rate using the recorded timestamps. To account for temporal jitter across streams, the multivariate time-series reconstruction recursively estimated the inter-packet intervals and fitted a linear time-series to match the N of samples per packets and across data packets (similar procedures described in ^91^). See paragraph 2.4.1 for further details.

### 2.3 Experimental Paradigm

The experiment consisted of three main blocks, each comprising a series of 7-minute tasks targeting distinct cognitive and behavioral states: (1) rest, (2) action; (3) perception. Tasks across the three main blocks were interleaved to prevent participants falling asleep.

Except for the spontaneous walking task, participants were seated comfortably in a quiet, dimly lit room and were instructed to maintain fixation at a cross displayed in front of them and minimize any movements. Task instructions were delivered verbally following a written protocol. The experimenter was seated in the same room as the participants but always remained outside their field of view and minimized acoustic contamination.

#### 2.3.1 Rest

The first block included eyes open resting-state and ‘interoceptive attention’ conditions. In the former, participants were simply instructed to maintain fixation at a cross placed in front of them and minimize movements, while in the latter condition we instructed participants to relax and pay attention to their bodily sensations. Participants were not required to perform any interoceptive task.

The resting state condition served to establish individual physiological reference profiles: endogenous cardiac, breathing and neural activity, as well spontaneous eye movement and pupil dilation at rest. The ‘interoceptive attention’ condition was expected to capture fluctuations in body-brain coupling dynamics in function of attention. Data from the interoceptive condition was not utilized for the main analyses presented here.

#### 2.3.2 Action

The action block comprised four 7-minute tasks requiring participants to generate spontaneous or adaptive rhythmic behavior. The spontaneous production conditions were designed to characterize the temporal properties of self-generated motor rhythms (spontaneous motor tempo) and their coupling to endogenous bodily and brain signals ^42,48,49^.

1. **Spontaneous tapping:** Participants were instructed to tap at a comfortable, self-chosen rate, without any external pacing. Participants were free to utilize their preferred index finger, and responses were recorded on a response button displayed on the custom-made app.
2. **Spontaneous walking:** Participants walked at their preferred, self-selected pace without external cues following a chosen trajectory inside the University campus.
3. **Spontaneous reading:** Participants read out loud the first chapter of the book titled ‘The tell-tale brain’ at their own pace. As for the tasks above, to maintain ecological validity, no temporal constraints were imposed on participants: they could pause and continue behaving freely.
4. **Adaptive tapping:** Participants were instructed to synchronize their tapping to a rhythmic equitone auditory sequence. The sequence comprized 7 isochronous blocks varying in their rates (.5, 1, 1.5, 2, 2.5, 3 Hz) presented in a random order (‘*randperm*’ in MATLAB) and with varying number of intervals (ranging from 20 to 40 intervals (‘*randi([20,40])*’)). The resulting sequence enabled participants to synchronize their tapping to predictable adjacent intervals and required them to speed up or slow down to tempo changes. Data from this task were not used here.

#### 2.3.3 Perception

The perception block consisted of four 7-minute passive listening conditions in which participants listened to auditory stimuli delivered via loudspeaker at a comfortable fixed volume (40% of the max output of the computer in use). While described here for completeness, data from the perception block was not used for the purpose of this manuscript.

1. **Isochronous sequence:** Participants listened to an isochronous auditory sequences comprising equitone sounds (F0 = 440 Hz, duration = 50 ms, 10ms rise and fall times) presented at a rate of 1.5Hz and generated at a sampling frequency of 14kHz. The task was designed to test neural entrainment to temporally predictable auditory sequences (^92,93^).
2. **Jittered sequence:** Participants listened to a temporally jittered auditory sequence comprising equitone sounds (F0 = 440 Hz, duration = 50 ms, 10ms rise and fall times). The inter-onset interval was derived from the speech sequence (listening task 3 below): the speech sequence underwent band-pass frequency filtering (4^th^ order, two-directional Butterworth filter); we extracted the amplitude envelope and determined sound onset timing based on an amplitude threshold criterion. The resulting jittered sequence had a mean rate of 3Hz (range 1-4Hz). The task was designed to test neural entrainment to rhythmic streams.
3. **Speech and Poems:** Participants listened to naturalistic speech excerpts recorded at a sampling rate of 14kHz. While speech features semi-periodic sound onsets, poetic meter is thought to introduce a more strongly periodic rhythmic structure ^94,95^.

#### 2.3.4 Interoceptive assessment

At the end of the experiment, participants completed the Multidimensional Assessment of Interoceptive Awareness, Version 2 (MAIA-2; ^74^). MAIA-2 is a self-report, 8-factor model state-trait questionnaire designed to measure multiple dimensions of interoception ^75,96^. Self-reported scores in the 37-item scale converge into the following factors: Noticing, Not-Distracting, Not-Worrying, Attention Regulation, Emotional Awareness, Self-Regulation, Body Listening, and Trust.

### 2.4 Data analysis

All analyses were implemented in MATLAB (version 2025a; MathWorks, Natick, MA, USA) using a combination of custom scripts and functions (GitHub link) and the FieldTrip toolbox ^97^.

#### 2.4.1 EEG preprocessing pipeline

The EEG preprocessing pipeline mainly comprised five sequential stages: (1) import of multimodal data; (2) detection and interpolation of missing data samples resulting from Bluetooth transmission failures; (3) motion artifact removal; (4) cardiac artifact removal; and (5) blink artifact removal and frequency filtering.

##### Import

A fundamental characteristic of BLE data transmission is that sensor samples are bundled into notifications (packets) prior to transmission, and the receiving device assigns a single reception timestamp to the entire packet irrespective of its size. To assign a physiologically accurate timestamp to each individual sample, a linear model was fitted to the packet anchor points — defined as the reception timestamp of each packet paired with the 0-based index of its last sample within the full sample sequence. This approach treats BLE reception time as a noisy but unbiased estimate of the true sample time for the final sample in each packet and uses the nominal sampling rate to backproject timestamps within the packet. Formally, for each stream a robust ordinary least-squares regression was computed as t (*k*) = *a* + *b·k*, where *k* is the zero-based sample index and the coefficients *a* and *b* are estimated from the packet anchor pairs {*k*_last, *t*_received}. The estimated inter-sample interval *b* provided a data-driven verification of the actual sampling frequency (EEG: 256Hz; ECG: 130Hz; ACC: 50 Hz). This linear reconstruction yielded a strictly monotonic, jitter-free time vector for each stream, with Bluetooth reception jitter of up to ±25 ms absorbed by the regression rather than propagated to individual sample times. The maximum residual between packet anchor timestamps and the linear fit was 25.7ms for EEG, consistent with known BLE scheduling latency under concurrent packet traffic. Following individual stream reconstruction, all streams were resampled onto a common uniform time grid at a target frequency of 256Hz to enable multivariate analysis. Resampling was performed using piecewise cubic Hermite interpolating polynomial (‘*pchip*’) interpolation ^98^, which preserves peak morphology and monotonicity of physiological waveforms. Because both the Muse and the PolarH10 are timestamped by the single tablet system clock, the reconstructed time vectors share a common reference frame without requiring external hardware triggers or post-hoc clock alignment.

##### Missing data samples

Intermittent Bluetooth disconnections might have introduced structured temporal gaps in which all EEG channels were simultaneously missing in some recordings. Missing samples were identified either by NaN values in the raw data stream or by an automated flat-line detection algorithm that flagged contiguous segments of near-zero variance (threshold: standard deviation < 10^−10^ μV, evaluated within a 100 ms sliding window). Because these gaps affected all channels simultaneously, cross-channel spatial interpolation (e.g., spherical spline interpolation of individual electrodes ^99^) was not applicable. Instead, gap segments were classified by duration and reconstructed using a tailored strategy depending on the temporal scale of the empty data intervals. Short gaps (≤ 50 ms) were reconstructed using shape-preserving piecewise cubic interpolation (‘pchip’). Pchip differs from conventional cubic spline interpolation ^98^ in that it enforces local monotonicity at each knot point. Medium-duration gaps (51–150 ms) were handled using a Rauch-Tung-Striebel (RTS) Kalman smoother (Rauch et al., 1965; ^100^). A forward Kalman filter pass was first applied to the concatenated sequence of [left context | gap | right context], treating gap samples as missing observations. A backward RTS smoother pass was then applied to the forward-filtered state estimates, yielding the optimal posterior state sequence — in the minimum mean-squared error sense — conditioned on all available observations from both flanks. The bidirectional exploitation of the data is a key advantage of the RTS smoother over other methods (e.g., one-sided autoregressive (AR) prediction for medium-duration gaps; see below). Longer gaps (151– 1000 ms) were reconstructed using a forward-backward AR blending procedure. An AR model of order *p* = 30 was estimated independently from the pre- and post-gap flanking segments (context window: 500 ms) using the Burg lattice algorithm (Burg, 1968). A forward prediction was generated by iterating the AR recursion from the terminal samples of the left-flank segment into the gap; an equivalent backward prediction was produced by applying the same procedure to the time-reversed right-flank segment and then reversing the result. The two predictions were combined using a raized-cosine (Hanning) taper, with blending weights transitioning smoothly from unity (forward) to zero (backward) across the gap duration. This ensures signal continuity and first-derivative continuity at both gap boundaries, while preserving the instantaneous spectral characteristics of the signal. Gaps exceeding 1000 ms were retained as NaN values and annotated as ‘bad’ segments. These segments were excluded from all subsequent analyses. The proportion and total duration of rejected segments were recorded per participant as a data quality metric.

##### Motion removal

Prior to motion removal, EEG data were high-pass filtered at 1Hz using a zero-phase FIR filter (Hamming window; filter order: 3 × ⌊f_s_ / f_HP_⌋, rounded to the nearest even integer), applied using the forward-backward (‘*filtfilt’*) procedure. This step removed DC offsets and slow drifts attributable to electrode drift, sweat artefacts, and electrode displacement, which would otherwise inflate the covariance between the EEG signal and the inertial reference signals, reducing the specificity of subsequent motion regression.

Motion-induced artifacts in wearable EEG may arise from multiple concurrent mechanisms including electrode-scalp relative displacement and impedance modulation, whose combined effect on the recorded signal is a non-linear and non-stationary function of head kinematics ^101,102^. This contrasts with the motion artifact model implicit in fMRI preprocessing, where rigid-body realignment parameters serve as nuisance regressors in a static general linear model (GLM; Friston et al., 1996). Applying a static GLM to wearable EEG motion artifacts is inappropriate because the transfer function between IMU signals and EEG contamination is time-varying: it changes as a function of electrode contact pressure, body orientation, and movement type. A fixed regression weight estimated from the full recording might therefore systematically underestimates artifact magnitude in high-movement epochs and overestimate it during rest, leaving structured residuals in both conditions. Here, motion artifact removal was therefore performed using a recursive least-squares (RLS) adaptive filter ^101,102^, which updates regression weights sample-by-sample according to the RLS recursion. At each time step *t*, the predicted artifact is the inner product of the current weight vector w(*t*−1) and the reference signal vector x(*t*), and the weight vector is updated by: w(*t*) = w(*t*−1) + k(*t*) e(*t*), where e(*t*) = y(*t*) − w(*t*−1)^T^ x(*t*) is the residual error and k(*t*) is the RLS gain vector, computed as k(*t*) = P(*t*−1) x(*t*) / [λ + x(*t*)^T^ P(*t*−1) x(*t*)]. The inverse covariance matrix P(*t*) is updated as P(*t*) = λ^−1^[P(*t*−1) − k(*t*) x(*t*)^T^ P(*t*−1)]. The forgetting factor λ governs the effective memory of the filter: at λ = 0.9995 and a sampling rate of 256 Hz, the filter has an effective memory of approximately 8s, chosen to balance tracking speed against noise sensitivity. The covariance matrix was initialized as P(0) = δ^−1^I with δ = 100, reflecting a weakly informative prior on initial weight magnitudes. The reference signal design matrix comprised 19 regressors per time point: the six raw IMU signals (three-axis linear acceleration and three-axis angular velocity), their first-order temporal derivatives (approximating electrode velocity relative to the scalp), their second-order temporal derivatives (approximating electrode acceleration), and a DC bias term. The inclusion of derivative regressors was motivated by the finding that scalp electrode artifact amplitudes are determined by the rate of change of electrode-scalp displacement as much as by displacement itself ^102^. All IMU signals were smoothed prior to differentiation using a 20-ms moving-average filter to suppress amplification of high-frequency noise in the derivative estimates. All reference signals were z-scored to unit variance prior to adaptive filtering, improving the numerical conditioning of the RLS covariance update. Each EEG channel was processed independently. A post-hoc sanity check was applied: if the RLS procedure increased the signal variance of any channel (indicative of over-subtraction or spurious correlation between reference and neural signal), the cleaned signal was discarded and the original channel retained for that segment.

##### Cardiac Artifact Removal

Cardiac artifacts in scalp EEG arise from two overlapping sources: direct volume-conducted contamination of the ECG signal and the ballistocardiographic artifact — a mechanically-induced artifact caused by pulsatile scalp displacement at each heartbeat ^103^. In the context of a wearable device without electromagnetic shielding, both sources are expected to be present simultaneously. Because both are phase-locked to the cardiac cycle, they were addressed in a unified framework using the Optimal Basis Set (OBS) method ^103,104^. R-peaks in the simultaneously acquired ECG signal were detected using a simplified Pan-Tompkins algorithm ^88^. The ECG was bandpass-filtered between 5 and 15 Hz to isolate the QRS energy band, differentiated using a five-point derivative filter (coefficients: [1, 2, 0, −2, −1] × f_s_/8), squared to produce a unipolar signal, and integrated with a 150-ms moving-window integrator to smooth the energy profile. R-peak candidates were identified as local maxima of the integrated signal exceeding a threshold of 3 standard deviations above the mean, subject to a 400-ms refractory period to enforce the physiological constraint on minimum inter-beat interval. Each candidate was subsequently refined to the nearest local maximum in the raw ECG signal within a ±50-ms search window. Heart rate and R-R interval statistics were computed for each participant as a detection quality control metric. For each EEG channel, the signal was epoched around each detected R-peak using a window of −300 to +500 ms. The epoch matrix was mean-subtracted on a per-epoch basis to remove heartbeat-to-heartbeat DC offsets. An orthogonal basis set was then constructed from the demeaned epoch matrix via singular value decomposition (SVD), retaining the leading *n* = 4 basis vectors. This constitutes the optimal basis in the sense that these vectors capture the maximum proportion of inter-epoch artifact variance with the minimum number of dimensions ^103^. The choice of *n* = 4 was motivated by two considerations: (i) the BCG waveform is known to exhibit significant morphological variability across heartbeats due to respiration, electrode movement, and blood pressure changes, requiring more than a single average template to model adequately; and (ii) increasing *n* beyond 4–6 risks absorbing genuine neural signal components that may be weakly phase-locked to the cardiac cycle, such as heartbeat-evoked potentials ^105^. For each epoch, the artifact contribution was estimated by projecting the EEG segment onto the basis set via ordinary least-squares and subtracting the resulting estimate from the raw signal. A Hanning taper was applied to prevent discontinuities at epoch boundaries.

##### Final steps

Residual high-amplitude, (non-)stationary burst artifacts (e.g., such as those produced by impulsive head movements) or semi-periodic eye blinks were suppressed using an artifact suppression algorithm ^93,106–108^. Noisy (>4*median absolute amplitude derivative) time-windows and their neighboring time-points (20 ms on each side) were interpolated using the same interpolation strategy as mentioned above. Following the artifact removal steps, EEG data were bandpass-filtered between 1 and 40 Hz using a zero-phase FIR filter (Hamming window; order: 3 × ⌊f_s_ / f_low_⌋, rounded to the nearest even integer; applied via forward-backward ‘*filtfilt’*). The 40-Hz upper cutoff was chosen to exclude residual line noise and high-frequency EMG contamination (∼50Hz) while preserving all frequency bands of interest for the present analyses. The lower 1-Hz cutoff was redundant with the preliminary high-pass stage but was retained to ensure filter roll-off symmetry and to suppress any slow drifts introduced by artifact removal steps.

The total proportion of data excluded (marked as ‘bad’ and not interpolated) was computed for each participant and stored for data quality inspection.

#### 2.4.2 Ocular data preprocessing pipeline

Raw pupil diameter and gaze position data exported from Tobii Pro Lab software version 25.7 were imported into MATLAB and processed following standard guidelines ^109–112^. The preprocessing pipeline mainly comprized three sequential stages: (1) assessment of data quality; (2) detection and marking of blink-related missing samples; (3) interpolation of blink-related missing intervals and low-pass filtering.

##### Data quality assessment

As wearable eye-tracking recordings can differ in quality (e.g., missing data) between the two eyes, a quality assessment was performed to determine whether binocular or monocular data should be retained for each recording. We considered two major criteria: percentage of missing samples relative to the entire recording; and left-right pupil diameter correlation. Missing samples were quantified separately for the left and right eyes. Recordings were excluded if both eyes contained more than 30% missing samples. If only one eye exceeded this threshold, the subsequent preprocessing relied on the contralateral eye. If both eyes contained less than 30% missing samples, we computed a left-right pupil correlation for paired valid samples (non-missing entries). We retained binocular data when the left-right correlation was >= 0.70; alternatively, the eye with fewer missing samples was preferred for next steps.

##### Blink detection

As blinks and brief eye tracking losses are commonly reflected in missing or invalid pupil samples ^111,112^, blink detection was based on transient missing segments in the pupil diameter time series. In monocular recordings, continuous missing segments lasting at least 100 ms were treated as blink candidates. In binocular recordings, missing segments were detected separately for the two eyes, and a blink was confirmed only when left- and right-eye missing periods overlapped or occurred within a 50 ms tolerance window. The confirmed blink interval was defined as the union of the corresponding left- and right-eye missing periods. Blink intervals separated by less than 250 ms were merged into a single event, and a 25 ms buffer was added to both sides of each merged interval to account for peri-blink signal loss. Samples within the final blink windows were set to NaN.

##### Interpolation and filtering

Following blink marking, NaN values in the pupil diameter time series, including blink-related and other transient missing samples, were reconstructed by interpolation. Interpolation was performed using the same adaptive method described in the ‘EEG preprocessing pipeline’ section. The resulting pupil data was then low-pass filtered at 2 Hz using an IIR filter to remove high-frequency noise while retaining slower pupil fluctuations ^109^. Finally, in binocular recordings we averaged the left- and right-eye pupil traces.

#### 2.4.3. Gait cycle reconstruction

Since accelerometer and gyroscope signals from MUSE data was often lost during recordings, gait cycle reconstruction was performed on the three-axis accelerometer data recorded by the Polar H10 chest band (see Suppl. Fig. 1). Data import followed the same procedure describe above.

As the sensor was not rigidly aligned to anatomical axes, a body-frame-independent measure was used: the signal vector magnitude (SVM), computed as the Euclidean norm of the three dynamic acceleration components after removing the mean of each axis across the full recording epoch to eliminate the static gravitational component ^113,114^. The resulting SVM time series — expressed in ms^−2^ following unit conversion from device-native *g* values — captures the net body acceleration irrespective of sensor orientation, and exhibits quasi-periodic oscillations locked to the gait cycle. The SVM was subsequently bandpass-filtered using a zero-phase fourth-order Butterworth filter with a passband of 0.5–3 Hz. The lower cutoff eliminates residual slow drift and postural sway, and the upper cutoff attenuates high-frequency mechanical vibrations; this passband encompasses the full physiological range of human walking and running cadences (approximately 30–180 steps per minute) ^115,116^. Prior to gait metrics computation, walking bouts were identified using a sliding-window variance thresholding approach applied to the unfiltered SVM. Consecutive two-second windows were classified as containing locomotion when the within-window SVM variance exceeded an empirically set threshold and classified as stationary otherwise. Contiguous locomotor windows were merged into single bouts. Gait analyses were conducted exclusively within identified walking bouts to avoid contaminating periodic metrics with data at rest. The primary gait periodicity metrics were derived from the normalized autocorrelation function (ACF) of the filtered SVM, following the methodology of Moe-Nilssen and Helbostad ^114^. The ACF was computed over lags from zero to four seconds. For a symmetric bipedal gait cycle, the ACF exhibits a first dominant peak at the step period (*T*_step; one peak per footfall) and a second dominant peak at the stride period (*T*_stride = 2 × *T*_step; one complete gait cycle). These peaks were identified using a prominence-constrained local maxima algorithm (minimum peak prominence: 0.05 normalized ACF units; minimum inter-peak distance: 0.25 s corresponding to a maximum cadence of 240 steps per minute). Step-by-step gait metrics were derived using local maxima detection on the bandpass-filtered, walking-masked SVM. Each local maximum was interpreted as corresponding to a heel strike event — the moment at which the center of mass reaches peak vertical displacement during single-limb support, which produces the dominant upward impulse detectable at the sternum ^113^. A minimum inter-peak interval of 0.25 s was enforced to suppress spurious detections, and a minimum peak prominence threshold of 10% of the signal maximum was applied to reject low-amplitude noise peaks. From the resulting step timestamp sequence, instantaneous cadence was computed as 60 divided by each consecutive inter-step interval (in seconds), yielding a time-resolved cadence series in steps per minute. The stride period was defined as the mean of every second inter-step interval (step pairs), and stride-to-stride variability was quantified as the coefficient of variation (CV; standard deviation / mean × 100%) of the stride period distribution, a metric with established clinical relevance as an index of gait regularity and fall risk ^50^. As an independent cross-check of the cadence estimate, a Hann-windowed discrete Fourier transform was applied to the walking-epoch SVM, and the frequency of the dominant spectral peak within the 0.5–3 Hz passband was identified. Agreement between the ACF-derived, peak-detection-derived, and FFT-derived cadence estimates within ±5 steps per minute was used as a quality criterion for each walking bout.

#### 2.4.4 Body-Brain-Behavioral rates

Two families of signals required different estimators: continuous signals (e.g., pupil diameter, respiration, speech) were analyzed in the frequency domain; event series (e.g., taps, saccades) were analyzed from the intervals between successive events. For every estimate we asked two questions: what is the predominant rate, and is there evidence that the signal is rhythmic at all? Where two independent estimators were available, we quantified their agreement. Details are provided below for each signal.

##### Body

Heart rates (HR) were estimated in each task and participant based on the inter-beat interval (IBI) in successive R-peaks (1/IBI). R-peaks in the ECG signal were detected using a simplified Pan-Tompkins algorithm (see details in ‘Cardiac artifact removal’ in paragraph 2.4.1). Continuous IBIs allowed estimating intra-individual HRs (Suppl. Fig. 2A) within and across tasks as the median HR. Group-level HRs at rest are reported in Fig. 2.

Breathing rates (BR) were estimated in each task and participant based on the inter-peak interval in successive inspiration peaks. Inspiration peaks in the EDR signal were detected using a peak detection algorithm. Intra-individual BRs and median BRs are reported in Suppl. Fig. 2B. Group-level BRs at rest are reported in Fig. 2.

Saccadic rates (SaR) were estimated utilizing Tobii’s gaze vector data. Gaze was converted to visual angle using the scene-camera field of view, and then to angular velocity between consecutive gaze points. Saccades were samples exceeding an adaptive threshold (median plus six robust standard deviations of speed, minimum 30°/s), with durations of 20–150 ms and amplitudes of at least 1° ^117^. Events within 50 ms of missing data (blinks) were discarded. Next, we calculated the SaR as described in ‘event series’ below. Importantly, this average saccadic rate is not representative of a truly endogenous, periodic rhythm. In fact, histograms of the inter-saccade intervals revealed a nearly uniform distribution over time, and across participants, indicating lack of a true rhythm. Consequently, SaRs will enter the BBB frequency architecture as a descriptive measure of the median saccadic rate over time, but do not represent an endogenous periodic signal. Group-level SaRs during walking are reported in Fig. 2. Pupil dilation rates (PDil) were estimated in each task and participant via fast Fourier transform. Continuous pupil dilation time-series were segmented in overlapping 10 s windows (1 s sliding), obtaining a frequency resolution of .1 Hz. Spectral power was calculated as the squared absolute value of the complex Fourier output. Individual and task-specific PDil were identified as the most prominent amplitude peak in the frequency range between .15 - .3Hz (given an hypothesized PDil >.1Hz ^118–120)^. Group-level data at rest is reported in Fig. 2 and Suppl. Fig. X.

##### Brain

Individual alpha frequency (IAF) peaks were obtained via fast Fourier transform. Continuous EEG signals from the two resting conditions were segmented in overlapping 10 s windows (1 s sliding), obtaining a frequency resolution of .1 Hz. Spectral power was calculated as the squared absolute value of the complex Fourier output. Frequency-domain data were normalized (z-scoring) and averaged across the two frontal channels. IAF peaks were identified as the most prominent amplitude peak in the frequency range between 8-12 Hz. Intra-individual and group-level IAF peaks are provided in Suppl. Fig. 2D (bottom right): the violin plot pools IAF peaks across moving windows and individuals in the Resting state condition. On its left, the Fourier spectrum obtained by averaging frequency-domain data across moving windows within and across participants, per condition. The individual median alpha peak in the rest conditions is provided in the insert in Suppl. Fig. 2D (left). IAF peaks across participants in the Resting state condition are provided in Fig. 2.

##### Behavior

Tapping rates (TR) were estimated in each task and participant based on the inter-onset interval in successive tap onsets. Group-level TRs at rest are reported in Fig. 2.

Walking rates (WR) were obtained from the walking task based on the inter-step intervals estimated via gait cycle analyses (see ‘Gait cycle reconstruction’ and Suppl. Fig. 1). Intra-individual WRs and group median WR are provided in Fig. 2.

Voice onset rates (VR) were estimated with the temporal modulation spectrum ^121^. The audio signal was split into 32 logarithmically spaced frequency bands (100–7000 Hz). The resulting amplitude envelope of each band was downsampled to 100 Hz before obtaining his spectrum. The spectrum of each envelope was computed in 10-s windows, normalised and averaged across bands. We defined the predominant speech rate as the modulation-spectrum peak in the rage of 1-10 Hz. To relate this spectral rate to discrete events, we also identified two event types in the broadband amplitude envelope: v*oice onsets* and s*yllable nuclei*. To obtain voice onsets, the speech signal underwent band-pass frequency filtering (4^th^ order, two-directional Butterworth filter) and a threshold criterion determined sound onset timing based on the amplitude envelope of the signal. The threshold was set per recording at 25% of the range between background (10th percentile) and loud speech (95th percentile) in decibels. Voice onsets rates were obtained as the inter-onset interval between successive energy peaks in the recorded speech sequences, spacing at least 50ms apart. Syllable nuclei were intensity peaks separated by dips of at least 2 dB and at least 100 ms apart ^122^. Syllable-nucleus rate was expressed per second of recording, comparable to the modulation spectrum, and per second of speech (articulation rate).

Importantly, these analyses were not specifically designed to isolate any language or speech specific component; in turn, we avoid referring directly to ‘syllable’ or ‘speaking rate’ and rather adopt the more general term ‘voice onset rates’. Intra-individual VRs are provided in Suppl. Fig. 3. Group-level VRs are reported in Fig. 2.

##### Continuous signals: spectral peaks

Physiological time series have power spectra that decline steeply with frequency (“1/f” or aperiodic background). The highest value in a frequency range is therefore often its lowest bin, which is not our rhythm of interest. We focused on the local spectral maximum rising above the aperiodic background. Power spectra were computed with Welch’s method (Hann-tapered, 50% overlapping windows, linear detrending per window; Welch 1967). The aperiodic component was fitted robustly in log–log space, following the logic of spectral parameterization ^123^. This approach was applied for pupil diameter, breathing, brain (Suppl. Fig. 2) and speech signals (Suppl. Fig. 3).

##### Event series: rates and rhythmicity

For each event series, we computed inter-event intervals and removed pauses (intervals longer than three times the median interval) so that interruptions did not count as slow cycles. The rate was defined as the inverse of the median interval, which is robust to occasional missed or extra events. Variability was summarized by the coefficient of variation. Rhythmicity was assessed with a windowed Rayleigh test: within sliding windows, we asked how strongly events cluster at a consistent phase of a candidate period, for each candidate frequency ^124^. Significance was assessed against surrogates in which each event was displaced randomly by up to half the median interval, thus destroying periodic timing while preserving slow changes in rate. Because any regular process (for example, a process with a refractory period) eventually beats this null with enough events, we also required a minimum phase-locking strength of 0.3 (0 = none, 1 = perfectly periodic). Rhythm ratios between successive intervals were inspected as an additional, assumption-free signature of isochrony ^125^.

This approach was employed for tapping, walking (Suppl. Fig. 4) and eye movement signals.

##### Agreement between estimators

Where a rhythm could be estimated by two independent methods (tap onsets vs tap-train spectrum (Suppl. Fig. 4); syllable nuclei vs modulation-spectrum peak (Suppl. Fig. 3)), agreement across participants was quantified on log-transformed rates. We report concordance correlation (CC; ^126^) and the intraclass correlation for absolute agreement (ICC; ^127^).

##### Distribution of individual rates within and across modalities

We asked two distinct questions: (i) How do individuals differ within each rhythm? And (ii) Does inter-individual variability scale with frequency?

i. For each modality, every participant contributes one rate, so observations are independent and identically distributed, rendering distributional tests valid. Two generative scenarios make different predictions. If an individual’s rate reflects a characteristic set point perturbed by many small, independent and *additive* influences, inter-individual differences should be approximately normally distributed around that set point (central limit theorem); this is expected for rhythms tightly constrained by a single dominant mechanism, such as biomechanical resonance or a homeostatic set point. If instead the rate results from the *multiplicative* combination of several independent factors (e.g., gains and time constants acting in series), its logarithm is a sum of independent terms and the rates should be lognormally distributed ^45^. A lognormal distribution implies right-skewed individual differences, variability that grows with the typical rate, and individual differences that are proportional rather than absolute, making the logarithmic scale the natural metric for comparing individuals. The two models converge when inter-individual variability is small (coefficient of variation (CV) below about 0.1), in which case they cannot be distinguished. For each modality we fitted normal and lognormal distributions to the raw rates by maximum likelihood; the lognormal likelihood included the Jacobian of the log transform, so the two fits were directly comparable. Both models have two parameters, and we compared them with the Akaike information criterion (ΔAIC = AIC_normal − AIC_lognormal; ΔAIC > 2 indicating support for the lognormal model and ΔAIC < −2 support for the normal model). We additionally report Lilliefors tests of normality on log-rates and the skewness of raw and log-rates.
ii. Buzsáki (2026) proposed that the variability of brain–body rhythms grows with their mean frequency. Under proportional (Weber-like) scaling, the standard deviation (SD) increases in proportion to the mean, so the CV is constant across rhythms. A variance proportional to the mean, as for counting processes, predicts SD ∝ mean^0.5; and identical absolute variability predicts no dependence on the mean. We regressed the log SD of individual rates on the log geometric mean rate across modalities. A slope of 1 indicates constant CV; a slope of 0.5 variance proportional to the mean; and a slope of 0 identical absolute variability. A 95% confidence interval for the slope was obtained by resampling participants (2,000 bootstrap samples).

##### Assembling the rate matrix

After characterizing the predominant rate across signals and tasks, we assembled a rate matrix per individual and across participants. While all signals displayed a predominant rhythm, saccades did not. Hence, they enter the rate matrix as a characteristic rate describing the median, but not an endogenous rhythm.

#### 2.4.5 MAIA-2

We processed data obtained from MAIA-2 following standard procedures ^74,128^ (also see: https://osher.ucsf.edu/research/maia), obtaining individual scores per participant across factors (see Suppl. Fig. 5). The MAIA-2 is a 37-item self-report questionnaire rated on a 6-point scale (0 = never, 5 = always). It yields eight scales: Noticing (awareness of body sensations; 4 items), Not-Distracting (tendency not to ignore or distract oneself from sensations of pain or discomfort; 6 items), Not-Worrying (tendency not to worry or experience emotional distress with sensations of pain or discomfort; 5 items), Attention Regulation (ability to sustain and control attention to body sensations; 7 items), Emotional Awareness (awareness of the connection between body sensations and emotional states; 5 items), Self-Regulation (ability to regulate distress by attention to body sensations; 4 items), Body Listening (actively listening to the body for insight; 3 items) and Trusting (experience of one’s body as safe and trustworthy; 3 items). Scale scores are the mean of their items, after reverse-scoring the negatively worded items of Not-Distracting and Not-Worrying. The MAIA-2 measures self-reported interoceptive sensibility, which is distinct from interoceptive accuracy as measured by behavioral tasks ^129^; results are interpreted accordingly.

Higher scores on Noticing, Attention Regulation, Emotional Awareness, Self Regulation, Body Listening, and Trusting may be interpreted as reflecting greater awareness of or attunement to bodily signals; higher scores on Not Distracting and Not Worrying may be interpreted as reflecting lower tendency to suppress or worry about body sensations.

#### 2.4.6 Frequency architecture

To test the hypothesis that a frequency architecture coordinates endogenous rhythms in the body, brain and behavior we adopted two complementary methodological approaches: (1) we statistically assessed the mathematical relationship between rhythms across modalities; (2) we tested whether individual rhythms could be predicted by a model utilizing information on inter-individual variation in other modalities. The predictive model was employed at the group level, as well as on two data-driven groups identified by clustering individuals depending on their interoceptive processing (as derived from MAIA-2). This step enabled testing the question of whether individual interoceptive levels modulate the relationship between body-brain-behavioral rhythms.

##### Rate relationship in BBB rhythms

After sorting body-brain-behavioral (BBB) rhythms in ascending order (Fig. 2B), we tested the hypothesis that BBB rhythms are characterized by a constant ratio between neighboring classes. On a logarithmic frequency axis, a constant ratio means equal steps: this would correspond to an arithmetic progression on a natural-logarithmic scale.

Different frameworks propose different ratios. Buzsáki ^2,19^ proposed a ratio of about *e*: 2.72; in this view, each rhythm class is roughly 2.7 times faster than the one below it. Starting at 0.25 Hz, such a ladder has rungs near 0.25, 0.68, 1.8, 5.0 and 13.6 Hz. As an alternative, Klimesch ^1^ proposes the golden mean (φ: 1.618) to define the bandwidth between adjacent rhythm classes. The golden mean, as well as the *e,* would produce rates at which rhythms are maximally decoupled: i.e., avoiding interference between independent processes.

##### Rate ratio fit

Modalities were ordered by their group-median rate, and this order was kept for all participants. For each participant, log-rates were regressed on position in this order. The slope, converted back to a ratio, is the factor by which each rhythm is faster than the previous one, on average. For example, a ratio of 2 would mean that each rhythm runs at twice the rate of the one before.

We compared five descriptions of each participant’s rates at consecutive positions. Three fixed ladders had the ratio set to *e* (slope rate: 2.72), φ (1.618), harmonic (2), with only the starting point free. A common empirical ratio was estimated from all other participants (leave-one-participant-out), so that the best-fitting ratio was not judged on the data it was derived from. Finally, an individual model let each participant have their own ratio.

For each model, we computed the root-mean-square deviation of the observed log-rates from the ladder and then compared models with the Bayesian information criterion (BIC), which rewards fit and penalizes free parameters. Group-level evidence was the summed BIC across participants, expressed relative to the individually fitted model. A difference above 10 is conventionally considered very strong evidence ^130^.

As some rhythm modalities clustered in rather close proximity to each other (e.g., cardiac, tapping and walking rates: 1.4-1.6Hz), we repeated the comparison creating rhythm ‘classes’. Classes were formed from modalities whose group medians lay within a factor of <.5 of each other.

##### Rate anchoring to HR

We tested Klimesch’s prediction according to which each person’s heart rate sets the scale for defining other rhythm classes. In this view, rhythm classes lie at heart rate multiplied or divided by powers of two. For example, breathing should lie near one quarter of heart rate (two octaves below), and alpha near eight times heart rate (three octaves above). For each participant and class, we expressed the class rate in octaves relative to that participant’s own heart rate. Thus, we defined frequency bands spanning from the domain center multiplied by φ/2 to the center multiplied by 2/φ, i.e. about ±0.31 octaves. A rhythm placed at random therefore falls inside a band with a probability of 61%. For each class, we compared the proportion of participants inside a band with this chance level (one-sided binomial test and one-sided Bayes factor ^131^ ; uniform prior on proportions above chance).

To test whether heart rate acts as an individual anchor, we compared the mean deviation from the nearest domain obtained with each participant’s own heart rate against three alternatives: heart rate shuffled across participants (5,000 permutations), a fixed population anchor of 1.25 Hz, and the group-median heart rate. If heart rate sets an individual scale, anchoring at one’s own heart rate must outperform anchoring at someone else’s.

To test whether inter-individual differences in HR propagate to other rhythms, as predicted by single-pacemaker accounts of internal timing, we modelled each rate as a power function of resting HR, f_m = c·HR^β, estimated as a linear regression on natural-log-transformed rates (ln f_m = a + β ln HR). Working on logarithmic scales renders the analysis independent of measurement units and of the absolute frequency of each rhythm, and makes the exponent β directly interpretable as a scaling law, as is customary in allometric analyses of biological rates ^132,133^. β = 0 indicates that a rhythm is independent of HR; β = 1 indicates proportional scaling, i.e. a ratio f_m/HR that is conserved across individuals, as expected if rhythms were rigidly locked to a common pacemaker; intermediate values indicate partial (allometric) scaling. For each modality, β was estimated by ordinary least squares, with 95% confidence intervals from 5,000 bootstrap resamples of participants. We compared three models with the Bayesian information criterion (BIC): independence (β fixed at 0), proportional scaling (β fixed at 1), and free scaling (β estimated). Posterior model probabilities were derived from BIC differences under equal prior odds ^134^. In addition, we quantified the evidence for an association between log-rates with a Bayes factor for Pearson’s correlation (uniform prior on ρ). We report BF01 (evidence for independence relative to association): values above 3 were interpreted as moderate and above 10 as strong evidence for the null hypothesis.

##### Prediction models

To test whether inter-individual variation in one modality is statistically predictable from inter-individual variation in the remaining modalities, we implemented two complementary approaches based on (1) cross-modal rate deviation scores; (2) ridge-regression and cross-validation.

Participants showing more than 5 missing entries across modalities were removed, leading to a final sample of 37 participants.

##### Cross-Modal Rate Deviation Score

To quantify how consistently each participant’s physiological, neural, and behavioural rates co-vary relative to the group, we constructed a log-ratio representation for each subject. For every pair of modalities (*i*, *j*) with non-missing values, the log-ratio was computed as:

log-ratio(*i, j*) = log(*x*׈) − log(*x*׈)

yielding a vector of up to *M*(*M* − 1)/2 = 28 values per participant. Log-ratios remove absolute scale differences between modalities — which ranged from 0.2 Hz (pupil dilation) to 11 Hz (alpha frequency) — and instead capture the relative structure of each individual’s rate configuration. A **deviation score** was then computed as the root-mean-square difference between each participant’s log-ratio profile and the group median profile, calculated across all pairs for which both modalities were non-missing. Smaller deviation scores indicate that a participant’s cross-modal rate structure conforms to the group-typical pattern; larger values indicate a more idiosyncratic profile.

##### Association between Interoceptive Awareness and Rate Deviation

To test whether individual differences in interoceptive awareness were associated with the degree of cross-modal rate consistency, we computed Spearman rank correlations between each of the eight MAIA subscales and the deviation score, restricted to participants with valid scores on both measures. Spearman’s ρ was selected for robustness to distributional non-normality and potential outliers, which are likely at the present sample size. Bootstrap 95% confidence intervals were computed for each correlation using 2,000 resamples with replacement ^135^. To control the family-wise false discovery rate across the eight simultaneous comparisons, p-values were adjusted using the Benjamini–Hochberg procedure ^136^.

To identify which MAIA subscales uniquely and independently predicted deviation score after accounting for inter-subscale correlations, we fitted a Ridge regression model ^137^ with all eight MAIA subscales as predictors and the deviation score as outcome, both standardized to zero mean and unit variance prior to fitting. Ridge regression was preferred over ordinary least squares given the modest sample size and the well-documented inter-correlations among MAIA subscales ^128,138^, which can inflate ordinary coefficient variance. The regularisation parameter λ was selected by leave-one-out cross-validation on the analytic sample. Standardized coefficients are reported, reflecting the direction and relative magnitude of each subscale’s unique association with rate deviation after partialling out shared variance.

##### Cross-Modal Prediction Model in MAIA-Defined Groups

To test whether interoceptive awareness moderated cross-modal rate predictability, participants were divided into high- and low-interoception subgroups using two independent grouping criteria: a median split on Body Listening — the MAIA subscale most directly relevant to attending to bodily rhythms — and a median split on the MAIA total score. Within each subgroup, a leave-one-modality-out prediction model was applied: for each target modality *m*, the remaining seven modalities served as predictors and the model was trained to predict the held-out target from inter-individual variation in the predictor set. Prior to modelling, each modality was subjected to a rank-based inverse normal transformation applied within each subgroup independently, mapping empirical ranks to standard normal quantiles via the Blom formula. This transformation makes no parametric assumption about the underlying distribution, is robust to outliers, and places all modalities on a common scale — a prerequisite for regularized regression. Ridge regression with λ = 1.0 was used as the prediction model, with missing predictor values imputed as zero (equivalent to column-mean imputation in rank-normalized space).

##### Inter-Individual Variability and Target Modality Selection

Before modelling, the coefficient of variation (CV = SD/mean, computed on the raw scale) was computed per modality to quantify the degree of inter-individual variability available for prediction. R² is mathematically defined as 1 − SS_res/SS_tot; when SS_tot is near zero — as occurs when a modality is near-constant across participants — any prediction error produces deeply negative R² irrespective of model performance. Modalities with CV < .10 were therefore retained as predictor variables but excluded as prediction targets. This threshold resulted in the exclusion of walking rate (CV = .070) and speaking rate (CV = .074) as targets, while retaining them as predictors. The six remaining targets — pupil dilation (CV = .248), breathing rate (CV = .118), heart rate (CV = .135), tapping rate (CV = 0549), saccadic rate (CV = .765), and alpha frequency (CV = .135) — exhibited sufficient inter-individual spread to support meaningful prediction. As alpha frequency was represented at discrete FFT frequency-resolution steps (∼.125 Hz, corresponding to the inverse of the EEG epoch length), it produced 17 unique values across participants; in turn, we suggest R² estimates for this modality to be interpreted with appropriate caution.

##### Statistical testing

Prediction accuracy was evaluated using Monte Carlo repeated random subsampling cross-validation with 1,000 iterations. At each iteration, participants with non-missing target values were randomly partitioned into 80% training and 20% test sets. Predictions were pooled across all iterations prior to computing the coefficient of determination R², providing a stable accuracy estimate that avoids the high variability of per-fold R² at small within-group sample sizes ^139^. R² = 0 indicates chance-level performance; negative values indicate the model performs worse than predicting the group mean. Modalities with a coefficient of variation below .10 in the full sample — walking rate and speaking rate — were retained as predictors but excluded as prediction targets, as their near-constant values across participants render R² mathematically unstable as an accuracy metric.

## Author Contributions

**A. Criscuolo:** conceptualization, data collection, methodology, writing, reviewing and editing, project administration. **T. Liu:** data collection, methodology, reviewing and editing. **M. Schwartze, S. Kotz:** conceptualization, reviewing and editing.

## Data and code availability

Data and code in use here will be made publicly available online upon publication.

